# Temporal dynamics of bacterial and fungal communities during the infection of Brassica rapa roots by the protist *Plasmodiophora brassicae*: The impact of a pathogen on the plant root and rhizosphere microbiota

**DOI:** 10.1101/410555

**Authors:** Lionel Lebreton, Anne-Yvonne Guillerm-Erckelboudt, Kévin Gazengel, Juliette Linglin, Morgane Ourry, Pascal Glory, Alain Sarniguet, Stéphanie Daval, Maria J. Manzanares-Dauleux, Christophe Mougel

## Abstract

The temporal dynamics of rhizosphere and root microbiota composition was compared between healthy and infected Chinese cabbage plants by the pathogen *Plasmodiophora brassicae*. When inoculated with *P. brassicae*, disease was measured at five sampling dates from early root hair infection to late gall development. The first symptoms of clubroot disease appeared 14 days after inoculation (DAI) and increased drastically between 14 and 35 DAI. The structure of microbial communities associated to rhizosphere soil and root from healthy and inoculated plants was characterized through high-throughput DNA sequencing of bacterial (16S) and fungal (18S) molecular markers and compared at each sampling date. In healthy plants, Proteobacteria and Bacteroidetes bacterial phyla dominated the rhizosphere and root microbiota of Chinese cabbage. Rhizosphere bacterial communities contained higher abundances of Actinobacteria and Firmicutes compared to the roots. Moreover, a drastic shift of fungal communities of healthy plants occurred between the two last sampling dates, especially in plant roots, where most of Ascomycota fungi dominated until they were replaced by a fungus assigned to the Chytridiomycota phylum. Parasitic invasion by *P. brassicae* disrupted the rhizosphere and root-associated community assembly at a late step during the root secondary cortical infection stage of clubroot disease. At this stage, *Flavisolibacter* and *Streptomyces* in the rhizosphere, and *Bacillus* in the roots, were drastically less abundant upon parasite invasion. Rhizosphere of plants colonized by *P. brassicae* was significantly more invaded by the Chytridiomycota fungus, which could reflect a mutualistic relationship in this compartment between these two microorganisms.

## Introduction

All plant tissues including roots [1,2], leaves [3,4] and seeds [5,6] are surrounded by a large diversity of microorganisms assembled in microbial communities or microbiota. These microbial assemblies represent a continuum of symbiosis with the plant ranging from parasitic to mutualistic interactions with complex microbe-microbe and microbe-plant interactions. Plant growth and health (including development, nutrition, physiology and defence) is influenced by these hosted complex microbial networks. Indeed, microbiota can stimulate seed germination and plant growth, help plants fight off disease, promote stress resistance, and influence plant fitness [7]. Thus, the plant microbiota extends the capacity of plants to adapt to their environment and contribute in shaping the plant phenotype.

Among these plant compartments, root and rhizosphere are the most studied habitats for microbial communities owing to their great potential for plant nutrition and health [1,8,9]. These microbial communities are mainly recruited by the plant from the soil [2,10,11] which is considered as the main microbial seed bank [12]. Many of these microorganisms including Archaea-and Eubacteria, fungi, and oomycetes live in the rhizosphere, defined as the narrow zone of soil that is influenced by root secretions [13,14]. Microbial community assemblies in the rhizosphere are governed by both abiotic and biotic factors. Soil properties, geographical location and Corine Land Cover in interaction with agronomical practices are the main factors that structure these communities [2,15,16]. Plant species and plant genotypes also determine to a lesser extent which members from the soil pool of microorganisms can grow and thrive in the rhizosphere [10,15,17,18]. Plants may modulate the rhizosphere microbiota to their benefit by selectively stimulating microorganisms showing traits that are beneficial to plant growth and health [9,19]. Rhizodeposits released by plant are known to account for variations of the diversity of microbial communities in the rhizosphere [20]. The modifications of the diversity of microbial communities are then expected to mirror variations of the composition of rhizodeposits. These rhizodeposits include both water-soluble exudates and more complex organic compounds resulting from dead cells sloughed off roots [21]. The proportion of photosynthates released in the rhizosphere and composition of the corresponding rhizodeposits have been shown to vary during the plant’s life cycle according to changes in plant physiology during the course of development and the level of symbiotic associations [22]. In addition, the genetic structure of bacterial and fungal communities was shown to change significantly during the development of *Medicago truncatula* in both vegetative and reproductive stages and the intensity of mutualistic symbiotic association with AM fungi and Rhizobia [23]. By extension, changes in microbial diversity and composition following plant-bioagressor interactions is often hypothesized to be based on modifications of the plant chemistry, such as plant exudates [24,25] or root metabolites [26].

Microorganisms that are able to penetrate and invade the plant root internal tissues form the endosphere or root microbiota. The roots of more than 80% of plants are colonized by arbuscular mycorrhizal fungi (AMF) and thus host symbiosis occurs with few dominant and well-known microorganisms. On the contrary, inside the Brassicaceae family, plants are believed not to form a strong symbiosis with few dominant microorganisms but hosts many types of microorganisms, including Archaea- and Eu-bacteria, fungi, and unicellular eukaryotes, such as algae and amoebae [27]. So far, only few studies focused on the composition, the dynamics and the ecological functions of these microorganisms during the plant growth. In contrast to the rhizosphere, the plant roots feature highly specific microbial communities [28]. The diversity of these endophyte communities is much lower than that estimated for microbial communities outside the root [10,11]. At the interface between the rhizosphere and the roots, the rhizoplane is often defined as a specific habitat of the rhizosphere because it is colonized by microorganisms that are firmly attached to the root surface. However, selective extraction and analysis of this compartment using culture-independent molecular methods and high-throughput sequencing are technically difficult and, consequently the role of the rhizoplane remains poorly understood [28]. Based on the composition of the rhizosphere and root microbiota, it has been proposed that the plants could assemble their microbiota in two steps, with the first one involving a rapid recruitment of microorganisms in the vicinity of the root and a second step being their entry inside the root [29]. However, the second step is more complex than the first one, with each root niche playing a selective role in microbiota assembly [15].

Even though the composition and recruitment mechanisms of these communities are being extensively investigated, only few studies dealt with the stability of these assemblies during the plant growth and under the effect of biotic stresses. Among biotic stresses, soil borne plant pathogens cause major economic losses in agricultural crops. Most of them are adapted to grow and survive in bulk soil but can also invade the root tissues to establish parasitic relationships with the plant. Since soil borne pathogens are already present in the soil before sowing, infections are usually early and occur during the vegetative stages of plant growth. To infect root tissues, pathogens have to compete with other microorganisms of the rhizosphere microbiota for available nutrients and microsites. One of the major roles of the rhizosphere microbial communities could to provide a frontline defence for plant roots against infection by soil borne pathogens [19]. This is valid not only for plant but also for animals and humans guts. Some of the mechanisms involved in the activity of these beneficial rhizosphere microorganisms are well studied and include several direct interactions with plant pathogens as well as indirect interactions via the plant by stimulating the plant immune system [30-31]. These mechanisms are well documented, using specific strains, for some rhizobacteria like *Pseudomonas* sp. and *Bacillus* sp., and for some fungi like *Trichoderma* sp. and non-pathogenic *Fusarium oxysporum*. However, most of the responsible microbial networks underlying these defences’ mechanisms are currently largely unknown. Recently, some metagenomic approaches provide us new opportunities to enrich our knowledge about the strong interactions between telluric pathogens and their living environment.

In this study, we described the dynamics of the root and rhizosphere (including the rhizoplane) communities of a Brassicaceae species during its vegetative stages, and we analysed the effects of a parasitic infection by *Plasmodiophora brassicae* use as a model system, on the composition and dynamics of these microbial communities. This pathogen is responsible for clubroot disease, a serious disease for many members of the Brassicacae family. *Plasmodiophora brassicae* Woronin is a soil borne obligate protist within the class Phytomyxea (plasmodiophorids) of the protist supergroup Rhizaria [33]. The pathogen life involves three stages: survival in the soil as resting spores, root hair primary infection and finally secondary cortical infection [34]. This process is accompanied by the hyperplasia and the hypertrophy of infected roots, resulting in formation of club-shape galls on the root. The tissue disruption associated with large clubs reduces nutrient and water transport within the plant, and consequently reduces plant growth and yield. *Brassica rapa* subsp. *Pekinensis* (Chinese cabbage) was chosen as the plant model of Brassicaceae because the full *P. brassicae* life cycle was easily achieved under controlled conditions in this species. We specifically addressed the following questions: (i) what is the dynamics of root and rhizosphere communities of Chinese cabbage during the vegetative stages of plant growth? (ii) How does *P. brassicae* affect the composition of bacterial and fungal rhizosphere and root communities at each of its life cycle stages? and (iii) which microbial species are selected following the infection by *P. brassicae*? To address these questions, a time-series experiment was conducted under controlled conditions. First, bacterial and fungal metagenomes of root and rhizosphere communities from non-inoculated (also called “healthy”) plants were described at successive time points. Then, the trajectories of microbial communities from healthy and inoculated plants cultivated in the same conditions were compared over time to analyse the effect of *P. brassicae* on the composition and stability of community assemblies in *B. rapa* plant roots.

## Materials and methods

### Materials

#### Soil

The experimental soil used for this study was collected at the INRA experimental site of La Gruche in Western Brittany (N: 48°08.44’, W: 01°47.98’). The topsoil (0-5 cm) was removed and the layer between −5 and −30 cm was harvested, homogenized, sieved at 4 mm and subsequently stored in 500 L containers at ambient temperature in the dark until further used. Physical and chemical properties of the soil were determined at the Arras soil analysis laboratory (F-62000, Arras, France). These properties were determined as: 13.3% sand, 70.9% silt, 15.8% clay, pH 6.2, 12.0 g.kg^-1^ of organic carbon, 1.2 g.kg^-1^ of mineral N and 20.8 g.kg^-1^ of organic matter.

#### Pathogen

The selection isolate eH used in this study belongs to the pathotype P1 [35], according to the host differential set established by [36]. This isolate was kindly provided by J Siemens (University of Dresden, Germany). It was propagated on Chinese cabbage as root galls, harvested, washed and stored at −20°C.

#### Plants

Seeds from the highly clubroot susceptible *Brassica rapa* spp. *pekinensis* cv. “Graanat” (ECD5) were used in this study to conduct the experiments. *B. napus* ssp. o*leifera* cv. “Nevin” (ECD6), *B. napus* ssp. *rapifera* cv. “Wilhelmsburger” (ECD10) and *B. napus* ssp. *oleifera* (Brutor), which constitute with *B. rapa* spp. *pekinensis* cv. “Graanat” (ECD5) the host differential was used as control to evaluate the infection success [36].

## Experimental design

### Plant growth assay and inoculation

ECD5 plants were cultivated in pots filled with 400 g of the experimental soil mixed with sterilized sand in the ratio 2:1. The experiment was conducted under a randomized complete block design using three blocks consisting of three replicates of four plants each. In each block, replicates were randomly distributed and placed in a greenhouse under the following conditions: 16 hours light (day) at 22°C and 8 hours dark (night) at 19°C. A mean photosynthetically active photon flux density of 150 μmol.m^-2^.s^-1^ at plant level during the 16 hours daylight was maintained. Some pots without plants were designated “bulk soil”.

Inoculum was prepared from three galls stored at −20°C as described previously [37]. In brief, spores were extracted by thawing the frozen galls at room temperature, and then homogenizing in 100 mL of sterilized water at high speed for 2 min. The resulting spore suspension was filtered through two sieves (250 and 100 μm pore diameters). The spore concentration was determined with a Malassez cell and adjusted to 1 × 10^7^ resting spores.mL^−1^. Ten-day-old seedlings were inoculated by pipetting 1 mL of spore suspension containing 1 × 10^7^ spores.mL^−1^ onto the soil surface at the base of each seedling. Non-inoculated plants and bulk soil were poured with sterile water. All pots, including bulk soil controls, were watered periodically every three days from the top with 8 mM Hoagland solution to maintain a water retention capacity between 70 - 100%.

### Quantification of plant traits

To follow the kinetics of plant growth, four plants per replicate were analyzed at 10 (T1), 17 (T2), 24 (T3), 33 (T4) and 37 (T5) days after sowing (DAS). Standard parameters were recorded: number of leaves per plant, shoot and root fresh weight, plant leaf areas, plant height and root length. Statistical analyses were performed using the R software [38]. Data were compared between healthy and diseased plants using linear models [LMM; function “lmer”, package “lme4”, [39]]. Pairwise comparisons of least square means (LSMeans) were performed using the function “lsmeans” [package “lsmeans”, [40]] and the false discovery rate (FDR) correction for p-values [41].

### Symptom development and clubroot severity measurement

Disease severity was assessed in inoculated plants during the vegetative stage of plant growth at 0 (T1), 7 (T2), 14 (T3), 23 (T4) and 35 (T5) days after inoculation (DAI) with *P. brassicae*, corresponding to 10, 17, 24, 33 and 45 DAS, respectively. Clubroot severity was recorded using the scale: 0, no visible swelling; 1, very slight swelling usually confined to lateral roots; 2, moderate swelling on lateral roots and taproot; 2+, severe clubs on all roots, but some roots remain present; and 3, no root left, only one big gall. A disease index (DI) was calculated as described by [42]: DI = (n1*25 + n2*50 + n2+*75 + n3*100)/N, where “ni” is the number of plants in the symptom class “i” and N is the total number of plants tested. Disease data were analyzed using a likelihood ratio test on a cumulative link model [40] [CLMM; function “clmm”, package “RVAideMemoire”, [43]]. Pairwise comparisons of LSMeans were then computed. To measure the hypertrophy of infected root, taproot width was also assessed at each date of sampling at 1 cm under the soil surface. Taproot width data were compared between non-inoculated (or healthy) and inoculated (or diseased) plants using a linear model [LMM; function “lmer”, package “lme4”]. Pairwise comparisons of LSMeans [function “lsmeans”, package “lsmeans”] and FDR correction for p-values were then performed.

### Sampling of “rhizosphere”, “root” and “bulk soil” compartments

Rhizosphere and root compartments from healthy and diseased plants were sampled at 10 (T1), 17 (T2), 24 (T3), 33 (T4) and 45 (T5) DAS. The “rhizosphere compartment” defined as the soil particles firmly attached to roots was collected by centrifugation of root washings. The “root compartment” was defined as the root tissues depleted of soil particles and epiphytic bacteria by sequential washing and sonication treatments and was therefore enriched in root-inhabiting bacteria.

Rhizosphere and root samples were collected from planted pots in a soil depth of −1 to −6 cm from the surface. Roots were separated from non-adhering soil particles, collected in 15 mL Falcon containing 20 mL sterile water and vortexed for 1 min. Seminal and nodal roots were included in the analysis. After vortexing, roots were transferred in a sterile Petri dish and subjected to a second washing treatment with 5 mL sterile water. Double washed roots were transferred in 5 mL sterile water and sonicated twice for 3 s at 40 Hz to detach microbes living in close association with root tissues. Roots were transferred in a Petri dish, cut into fragments smaller than 5 mm, ground to a powder with a pestle in liquid nitrogen-chilled mortar with Fontainebleau sand and stored at −80°C until further analysis. The soil suspensions collected in the Falcon tubes or in the Petri dishes after the first, the second washing treatments and the sonicated solution were pooled, centrifuged at 4,000 g for 20 min and the pellet, referred to as the rhizosphere, was frozen in liquid nitrogen and stored at −80°C until further analysis.

Soil samples were collected from unplanted pots at T1, T3 and T5 in a soil depth of −1 to −6 cm from the surface. The soils from four pots were pooled, transferred in 10 mL sterile water and vortexed for 1 min. The soil suspension was centrifuged at 4,000 g for 20 min and the pellet, referred as the bulk soil, was frozen in liquid nitrogen and stored at −80°C until further analysis.

## DNA extraction and pathogen quantification

### Root and soil DNA extraction

The GnS-GII protocol was used for root and DNA extraction [44]. For root samples, five to 150 mg of each root sample were homogenized for 3 × 30 s at 4 m.sec^-1^ in a FastPrep®-24 (MP-Biomedicals, NY, USA) in 2 mL of the “lysing matrix E” solution from MpBio containing 100 mM Tris (pH 8.0), 100 mM EDTA (pH 8.0), 100 mM NaCl, and 2% (wt/vol) sodium dodecyl sulphate. The samples were incubated for 30 min at 70°C, and then centrifuged at 7,000 g for 1 min at 20°C. To remove proteins from the extracts, 1 mL of the collected supernatant was incubated for 10 min on ice with 1/10 volume of 3 M potassium acetate (pH 5.5) and centrifuged at 14,000 g for 5 min at 4°C. Finally, after precipitation with 900 μL of ice-cold isopropanol, the nucleic acids were washed with 70% ice-cold ethanol and DNA was resuspended in 200 μL ultrapure sterile water. DNA was separated from the residual impurities, particularly humic substances, by centrifuging through two types of minicolumns. Firstly, aliquots (100 μL) of crude DNA extract were first loaded onto Microbiospin (Biorad, Hercules, California, USA) columns of PVPP (PolyVinyl PolyPyrrolydone) and centrifuged at 1,000 g for 2 min at 10°C. Secondly, the eluate was purified with the Geneclean turbo kit (Q-Biogene, Illkirch, France). DNA concentration and purity were determined with a Nanodrop (Agilent).

The same protocol was used to extract DNA from soil samples except that, before homogenization in the FastPrep ®-24, 2 g of each soil sample were mixed with 5 mL of a solution containing 100 mM Tris (pH 8.0), 100 mM EDTA (pH 8.0), 100 mM NaCl, and 2% (wt/vol) sodium dodecyl sulphate in a 15 mL “lysing matrix E” Falcon tube from MpBio.

### Measurement of pathogen DNA amount in roots by real-time qPCR

Plant root colonization by *P. brassicae* was also monitored by quantitative PCR. The predicted 18S gene was used to estimate *P. brassicae* DNA amount per ng of total extracted DNA. A portion of this gene sizing 164 bp was amplified with the primers PbK1F/PbK1R (5’-TTGGGTAATTTGCGCGCCTG-3’/5’-CAGCGGCAGGTCATTCAACA-3’). All reactions were performed in 20 μL qPCR reaction using 10 μL of SYBR Green Master Mix (Roche), 1 μL of DNA (2.5 ng) and 0.08 μL of each primer (100 μM). Amplification conditions were as follows: 5 min at 95°C, followed by 45 two-step cycles at 95°C (10s) and 60°C (40s). Standard curves were constructed using serial dilutions of *P. brassicae* DNA extracted from resting spores. A linear model [LMM; function “lmer”, package “lme4”] was used to analyze the pathogen DNA quantification data. Pairwise comparisons of LSMeans [function “lsmeans”, package “lsmeans”] and FDR correction for p-values were then performed.

## Bacterial and fungal community composition and diversity

### Sequencing of 16S and 18S rDNA genes

The structure of microbial communities associated to soil and root samples collected during the experiments was assessed though amplification and subsequent sequencing of bacterial (16S) and fungal (18S) rDNA genes. PCR amplification and sequencing were performed at GenoScreen (Lille, France) using the Illumina Miseq platform to a 2 × 300 bases paired-end version with an adequate read assembly method. For soil and root DNA extracts, a 420 bp fragment of the V5-V7 region of the bacterial 16S rDNA gene was amplified using the universal bacterial primers 799F_16S (5’-AACMGGATTAGATACCCKG-3’) and 1223R_16S (5’-CCATTGTAGTACGTGTGTA-3’) [45,46]. Before sequencing, PCR products were purified to eliminate a 760 bp fragment corresponding to plant mitochondrial DNA amplification. A 530bp fragment of the fungal 18S rDNA that includes the variable regions V4 (partial) and V5 was also amplified using the primer pair NS22B (5’-AATTAAGCAGACAAATCACT-3’) and SSU0817 (5’-TTAGCATGGAATAATRRAATAGGA-3’) [47,48].

### Analysis of MiSeq sequencing data

After reads assembly, sequences were processed with GnS-PIPE bioinformatics platform developed by GenoSol platform and optimized for amplicons analysis [49,50]. The reads were filtered and eliminated if they harbored one or more ambiguities (Ns) or an average quality score below 30. A PERL program was applied to obtain strict dereplication (i.e. clustering of strictly identical sequences). After this initial quality filtering step, the reads were aligned with INFERNAL alignments [51] and clustered at 97% sequence similarity into operational taxonomic units (OTU) using another PERL program. All single-singletons (reads detected only once and not clustered) were then deleted in order to eliminate PCR chimeras and large sequencing errors. These final sequences were used to produce rarefaction curves. The retained high-quality reads were used for taxonomy-based analysis of each OTU using similarity approaches against dedicated reference databases from SILVA [52]. The raw data sets are available on the European Nucleotide Archive database system under the project accession number PRJEB26948. Root and soil samples accession numbers range from ERS2513216 to ERS2513353 for 16S and 18S rDNA.

### Alpha diversity

To compare bacterial or fungal composition among bulk soil, rhizosphere soil and root from healthy and diseased plants, the richness was characterized by the number of OTUs found in each sample. As metric of taxonomy diversity, Shannon diversity was also determined using the “vegan” package in R, version 2.2-1 [53]. Since values were conformed to normality assumptions, two-way Anova and *post-hoc* Tukey’s HSD test were used to examine pairwise differences between samples for these measures.

### Beta diversity

After normalization by sample size, OTU counts without at least a mean of one read per sample were removed from the analysis. The genera OTU counts were also rarefied to 1,000 counts per sample and Log2-transformed rarefied values were used to calculate a Bray-Curtis distance dissimilarity matrix using the function “vegdist” of the R package “Vegan”. The beta diversity distance matrices were plotted using a bi-dimensional Principal Coordinates Analysis (PCoA) using the function “plot” of the R package “Vegan”. To quantify the influence of each factor on the beta diversity, a canonical analysis of principal coordinates (CAP, [54]) followed by a permutation-based ANOVA (PERMANOVA) was performed using the R package “vegan” according to the method described by [55].

### Statistical analysis on phyla counts

To identify phyla enriched in rhizosphere and root microhabitats compared to unplanted soil and to compare phyla composition between samples collected from healthy and diseased plants, we employed linear statistics on Relative Abundances (RA) values (log2 > 5‰ threshold) using a script developed from the R package “Limma” [17]. Differentially abundant phyla between two samples were calculated using moderated t-tests. The resulting p-values were adjusted for multiple hypotheses testing using the Benjamini-Hochberg (BH) correction.

### Detection of differentially enriched OTUs

EdgeR is a workflow largely based on the free open-source R language and Bioconductor software [56]. This workflow was originally used to analyze count-based differential expression of RNA sequencing as part of transcriptome studies [57] and was recently adapted to metagenomic data analysis [26]. OTU counts without at least a mean of one read per sample were removed from the analysis. To normalize the data for each sample OTU count, the trimmed mean of M values normalization method (TMM) was used according to the method described by [58]. A Log2-transformation was performed on the normalized data for statistical comparisons. Threshold, normalization and transformation steps were performed using a custom R script. To identify differentially abundant genus in bacterial and fungal communities between sampling dates and treatments (non-inoculated or inoculated) in root or soil samples, EdgeR was used to fit a model with treatment (non-inoculated or inoculated) * sampling date (T1 to T5) terms to the count data in each compartment by using glmFit and glmLRT with tagwise dispersion and to test for significant effects of each term. EdgeR employs statistical methods supported on negative binomial distribution as a model for count variability. Data from root and rhizosphere soil were not analyzed together because composition biases between samples from these two compartments were not eliminated by TMM normalization. To examine whether having a diverged or conserved communities composition was associated with treatment * time effect, the model was fitted to subsets of the normalized counts data and used “contrasts” to identify genera with significant differential abundances in pairwise comparisons. A likelihood ratio test (LRT) was performed to specify the difference of interest and the resulting p-values were adjusted for multiple hypotheses testing using the Benjamini-Hochberg (BH) correction.

## Results

Chinese cabbage plants were cultivated in a greenhouse for 45 days. Ten days after sowing, the plants were inoculated or not with *P. brassicae*. The roots and rhizosphere soils from healthy (or non-inoculated) and diseased (or inoculated) plants were both sampled at 0 (T1), 7 (T2), 14 (T3), 23 (T4) and 35 (T5) days after inoculation (DAI) with *P. brassicae*, corresponding to 10, 17, 24, 33 and 45 days after sowing (DAS). Bulk soil was also sampled within non-cultivated plots at T1, T3 and T5. Microbial composition from each compartment and at each date of sampling was assessed through 16S and 18S high-throughput sequencing. No clubroot symptoms were observed and no *P. brassicae* DNA was detected in non-inoculated plants.

### Comparison of communities from root, rhizosphere and bulk soils in healthy plants: the rhizosphere effect

In the samples collected at T1, T3 and T5 from healthy plants and bulk soil, the greatest numbers of bacterial/fungal OTUs were detected in bulk and rhizosphere soils (2,240/1,242 and 2,280/1,220 OTUs on average, respectively) and a significant reduction of richness was observed in root compartment (530/677 OTUs on average) (S1 and S2 Figs). A significant reduction of bacterial and fungal diversities in the root samples compared to bulk and rhizosphere soils was also observed at each sampling date (S1 and S2 Figs). A temporal effect on bacterial richness and diversity was measured but only in the root compartment where the number of OTUs and the Shannon index were higher at T3. In each compartment, no temporal variations of fungal richness and diversity was measured.

When looking at the microbial composition, we found that root bacterial and fungal communities were clearly distinct from rhizosphere and bulk soil communities at each sampling date (S3 Fig). A canonical analysis constrained by the variables of interest revealed that for bacterial communities, the compartment explained 52.5% of the variance (p = 0.001; 95% confidence interval (CI) = 24.5%, 86.7%) and the sampling date explained 5.5% of the variance (p = 0.001, 95% CI = 4.7%, 6.4%) (S4 Fig). For fungal communities, the compartment explained 29.8% of the variance (p = 0.001, 95% CI = 18.7%, 49.4%) and the sampling date 11.8% of the variance (p = 0.001, 95% CI = 8.7%, 16.2%) (S4 Fig). Consistently, we observed at T5 a clear separation between root microhabitat and soil samples followed by segregation of the rhizosphere and bulk soil samples. To explain the variance observed, the significant effect of the sampling date was weaker than the compartment.

### Composition and dynamics of healthy Chinese cabbage rhizosphere and root microbiota

#### In the rhizosphere of healthy plants

In the rhizosphere of healthy plants, the most heavily-sequenced bacterial phyla found were Proteobacteria, Firmicutes, Actinobacteria and Bacteroidetes, with 86% to 90% abundances at each sampling date between T1 and T5 (Fig 1). Within the rhizosphere-inhabiting Proteobacteria, the α-Proteobacteria were over-represented compared to the β-, γ- and δ-Proteobacteria (Fig 1). Between T1 and T5, a significant increase of Proteobacteria (α and γ) and a decrease of Firmicutes were measured while no temporal variation of bulk soil composition at phylum level was observed (data not shown). At T5, the enrichment of members from the Proteobacteria and Bacteroidetes phyla significantly discriminated rhizosphere from bulk soil samples (data not shown). We tried to narrow down the bacterial community to those OTUs (≥ 97% sequence similarity), which showed a minimum relative abundance of 0.1% at least in one of the rhizosphere samples. A total of 429 OTUs were identified in the rhizosphere of healthy plants (S1 Table). At the genus level, OTU1 assigned as *Bacillus* (Firmicutes) dominated these rhizosphere communities at each sampling date with 12 to 18% relative abundances between T1 and T5. OTU4 (*Sphingomonas*, α-Proteobacteria), OTU7 (*Pseudolabrys*, α-Proteobacteria), OTU9 (*Sporosarcina*, Firmicutes), OTU6 (*Bradyrhizobium*, α-Proteobacteria), and OTU10 (*Rhodopseudomonas*, α-Proteobacteria) were also highly represented (S1 Table). No temporal variation of these dominant OTUs was observed between T1 and T5. However, several minor OTUs with significant relative abundance variations between two sampling dates were detected in these bacterial communities (Table 1).

**Table 1.**
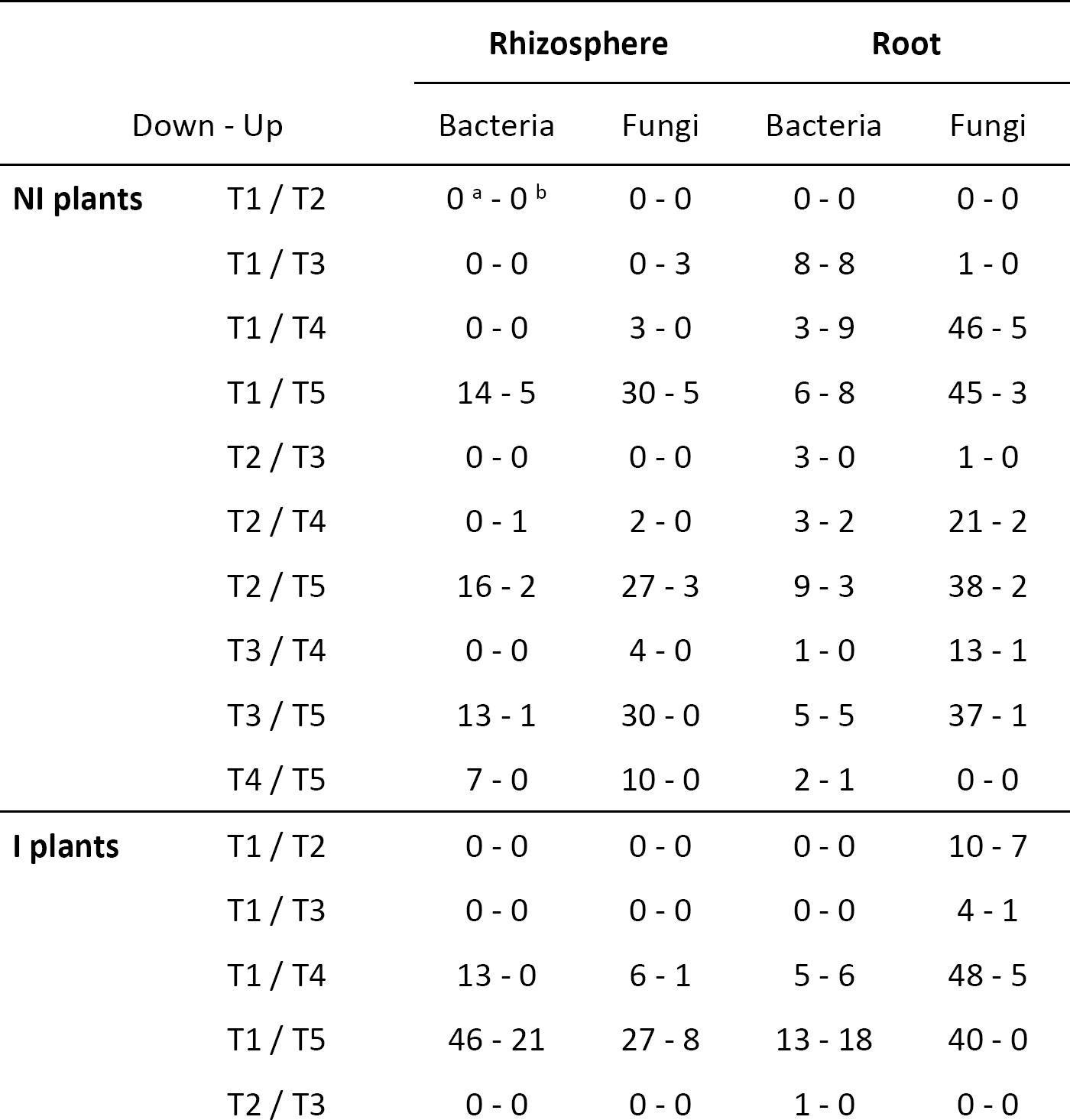
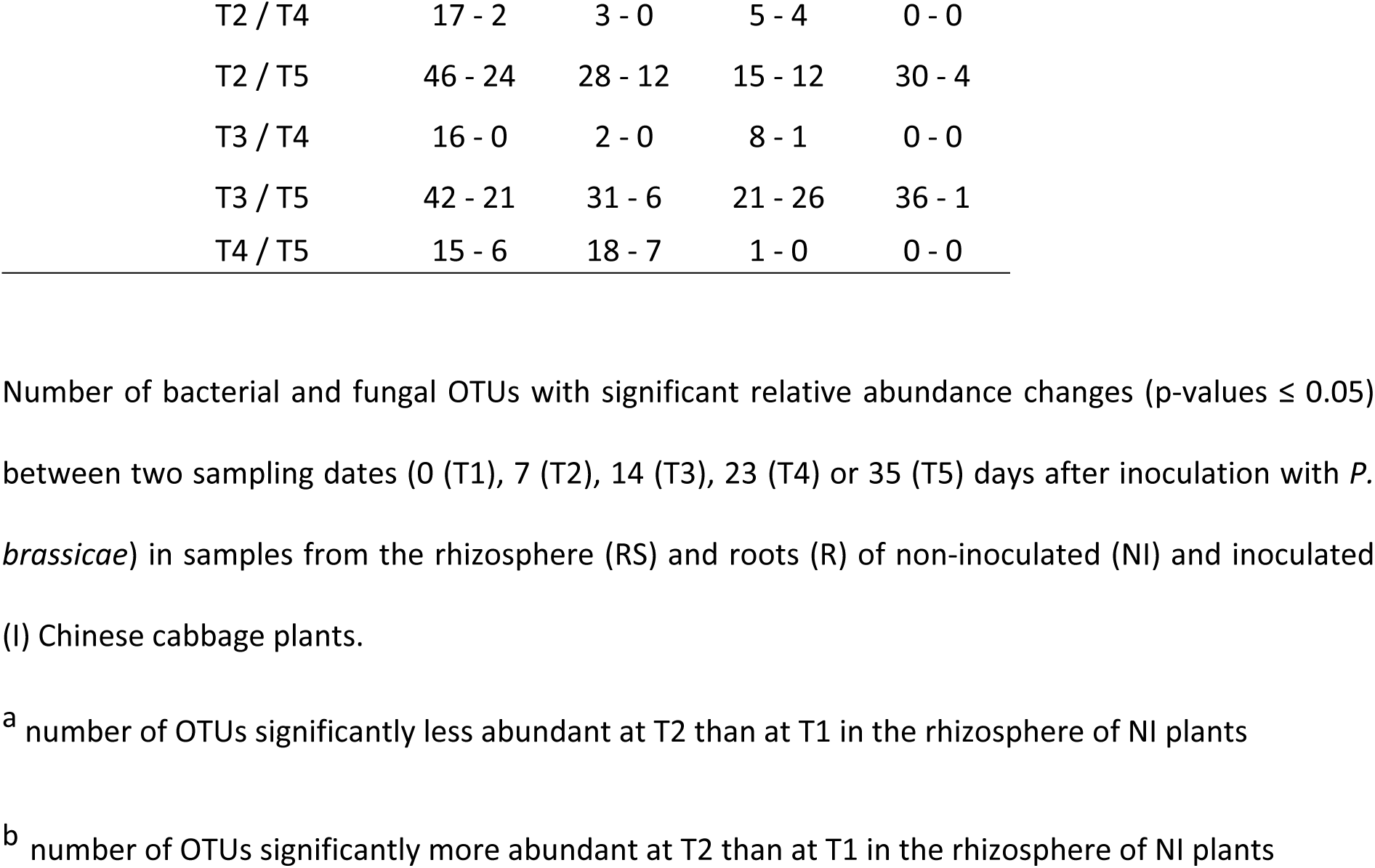
Number of OTUs with significant relative abundance changes between two sampling dates in the rhizosphere and roots of healthy (non-inoculated) and diseased (inoculated) plants.

|  |  | Rhizosphere |  | Root |  |
| --- | --- | --- | --- | --- | --- |
|  | Down - Up | Bacteria | Fungi | Bacteria | Fungi |
| **NI plants** | T1 / T2 | 0<sup>a</sup> - 0<sup>b</sup> | 0 - 0 | 0 - 0 | 0 - 0 |
|  | T1 / T3 | 0 - 0 | 0 - 3 | 8 - 8 | 1 - 0 |
|  | T1 / T4 | 0 - 0 | 3 - 0 | 3 - 9 | 46 - 5 |
|  | T1 / T5 | 14 - 5 | 30 - 5 | 6 - 8 | 45 - 3 |
|  | T2 / T3 | 0 - 0 | 0 - 0 | 3 - 0 | 1 - 0 |
|  | T2 / T4 | 0 - 1 | 2 - 0 | 3 - 2 | 21 - 2 |
|  | T2 / T5 | 16 - 2 | 27 - 3 | 9 - 3 | 38 - 2 |
|  | T3 / T4 | 0 - 0 | 4 - 0 | 1 - 0 | 13 - 1 |
|  | T3 / T5 | 13 - 1 | 30 - 0 | 5 - 5 | 37 - 1 |
|  | T4 / T5 | 7 - 0 | 10 - 0 | 2 - 1 | 0 - 0 |
| **I plants** | T1 / T2 | 0 - 0 | 0 - 0 | 0 - 0 | 10 - 7 |
|  | T1 / T3 | 0 - 0 | 0 - 0 | 0 - 0 | 4 - 1 |
|  | T1 / T4 | 13 - 0 | 6 - 1 | 5 - 6 | 48 - 5 |
|  | T1 / T5 | 46 - 21 | 27 - 8 | 13 - 18 | 40 - 0 |
|  | T2 / T3 | 0 - 0 | 0 - 0 | 1 - 0 | 0 - 0 |

|  |  |  |  |  |
| --- | --- | --- | --- | --- |
| T2 / T4 | 17 - 2 | 3 - 0 | 5 - 4 | 0 - 0 |
| T2 / T5 | 46 - 24 | 28 - 12 | 15 - 12 | 30 - 4 |
| T3 / T4 | 16 - 0 | 2 - 0 | 8 - 1 | 0 - 0 |
| T3 / T5 | 42 - 21 | 31 - 6 | 21 - 26 | 36 - 1 |
| T4 / T5 | 15 - 6 | 18 - 7 | 1 - 0 | 0 - 0 |

Number of bacterial and fungal OTUs with significant relative abundance changes (p-values $\leq 0.05$ ) between two sampling dates (0 (T1), 7 (T2), 14 (T3), 23 (T4) or 35 (T5) days after inoculation with *P. brassicae*) in samples from the rhizosphere (RS) and roots (R) of non-inoculated (NI) and inoculated (I) Chinese cabbage plants.
<sup>a</sup> number of OTUs significantly less abundant at T2 than at T1 in the rhizosphere of NI plants
<sup>b</sup> number of OTUs significantly more abundant at T2 than at T1 in the rhizosphere of NI plants

**Fig 1.**
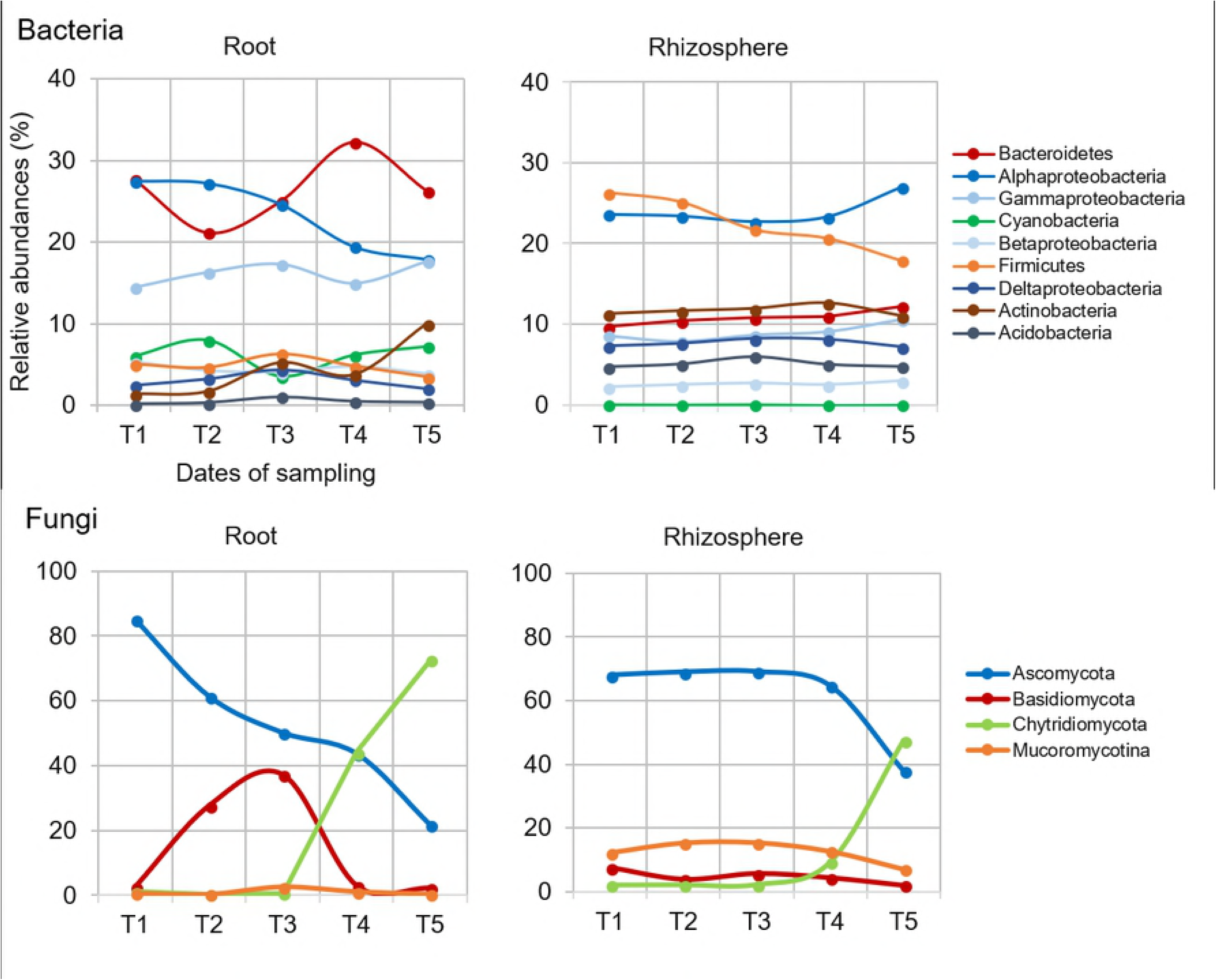
Temporal dynamics of the most abundant phyla-subphyla in bacterial (A) and fungal (B) communities from roots and rhizosphere of healthy plants. Mean values of abundance (expressed in %) were obtained from three replicates per condition and sampling date. Sampling dates refer to 10 (T1), 17 (T2), 24 (T3), 33 (T4) and 45 (T5) days after sowing. Phyla with relative abundances below 1% were grouped as “others”. In bacterial communities, the Proteobacteria phylum was divided into four subphyla: α-, β-, γ- and δ-Proteobacteria.

The fungal rhizosphere communities from healthy plants were largely dominated by Ascomycota, with 64% to 69% relative abundances between T1 and T4 (Fig 1). Three other phyla, Mucoromycotina (12% to 16%), Basidiomycota (3% to 8%) and Chytridiomycota (2% to 10%) were also detected but at lower frequencies. Until T4, the proportions of these four phyla compared to the whole fungal microbiota were relatively stable. From the 168 fungal OTUs identified with at least a relative abundance of 0.1% in one of these rhizosphere samples, OTU2 and OTU4, assigned as two Sordariomycetes, were detected at high frequencies between T1 and T4, but no temporal variations of these dominant OTUs were observed. At T5, fungi from the Chytridiomycota phylum were drastically more abundant (47.5%) than at the beginning of the kinetics and a decrease of Ascomycota (64.6% at T4 to 37.9% at T5) was measured (Fig 1). At this date of sampling, OTU1 and OTU16, assigned as two Chytridiomycota, were the most abundant OTUs with 18.3 and 16.4% relative abundances, respectively (S1 Table). Variations of other less dominant OTUs were also observed between T1 and T5 (Table 1). While the proportion of Chytridiomycota fungi strongly increased in the rhizosphere compartment at T5, their relative abundances remained low in bulk soil samples during all the kinetics.

#### Inside the roots of healthy plants

In the roots of healthy plants, bacterial communities were also dominated by Proteobacteria (41% to 51% RA between T1 and T5) and Bacteroidetes (21% to 33%) as in the rhizosphere (Fig 1). They also contained Actinobacteria (1% to 10%) and Firmicutes (3% to 7%), although to a lesser extent than in the rhizosphere samples. In the root samples, the α- and γ-Proteobacteria were over-represented compared to the β- and δ-Proteobacteria. Cyanobacteria were also detected. Between T1 and T5, more important variations of phylum frequencies occurred in the roots of healthy plants than in their rhizosphere (Fig 1). Actinobacteria increased significantly in frequencies while Proteobacteria decreased. A total of 202 genera were identified in these communities. The dominant OTUs were OTU2 assigned to a *Flavisolibacter* (Bacteroidetes) which relative abundances varied between 8% and 17%, OTU3 assigned to an unknown Cyanobacterium, OTU19 (*Devosia*, α-Proteobacteria), OTU12 (*Pseudomonas*, γ-Proteobacteria) and OTU5 (*Flavobacterium*, Bacteroidetes) (S1 Table). While the proportion of OTU2 strongly increased in the rhizosphere compartment of healthy plants between T1 and T5, the proportion of OTU19 decreased from 4.2% to 1.0% and no temporal variations was observed for the other main OTUs. Several other bacterial OTUs with significant variations in abundances from one date of sampling date to another were also detected in healthy plant roots (Table 1).

The root fungal communities were dominated by Ascomycota (85.1%) at T1 and replaced progressively by fungi from the Chytridiomycota phylum during the kinetics of plant growth. At T5, OTU1 which was assigned to the Chytridiomycota was detected in the roots of healthy plants at a very high frequency with a mean of 53% relative abundance (S1 Table). Variations of several minor OTUs were also observed in the fungal communities of diseased plants during the time-series experiment (Table 1).

To conclude, weak fluctuations were measured in the composition of rhizosphere and root communities of healthy plants before T4, whereas important changes occurred in these communities between T4 and T5. According to the variations of OTU relative abundances, these changes were first observed in the bacterial and fungal communities of plant roots and then in the rhizosphere (Table 1).

### Symptom development and clubroot severity

Differences of taproot width between healthy and diseased plants appeared at T3 and increased drastically between T3 and T5 (Fig 2). Disease index was low at T3 (DI = 16.7%), increased rapidly to 68.5% at T4 and reached a maximum of 86% at the end of the experiment (Fig 2). The amount of *P. brassicae* DNA followed a similar evolution profile (Fig 2). However, although DI increased between T4 and T5, there were no significant variations of *P. brassicae* DNA amount in roots at these time-points. Throughout the time of the experiment, no differences of leaf number and leaf area, plant height and shoot biomass were observed between healthy and diseased plants. In contrast, root length and biomass of inoculated plants decreased significantly but only between T4 and T5 (S2 Table). At the end of the experiment, some galls had become brownish and some mature resting spores were observed in gall tissues. According to these results, the duration of the life cycle of *P. brassicae* infection in Chinese cabbage was approximately 35 days in our experimental conditions. The root hair infection and the beginning of the cortical infection stages occurred before 14 DAI, and clubroots formed gradually between 14 and 35 DAI.

**Fig 2.**
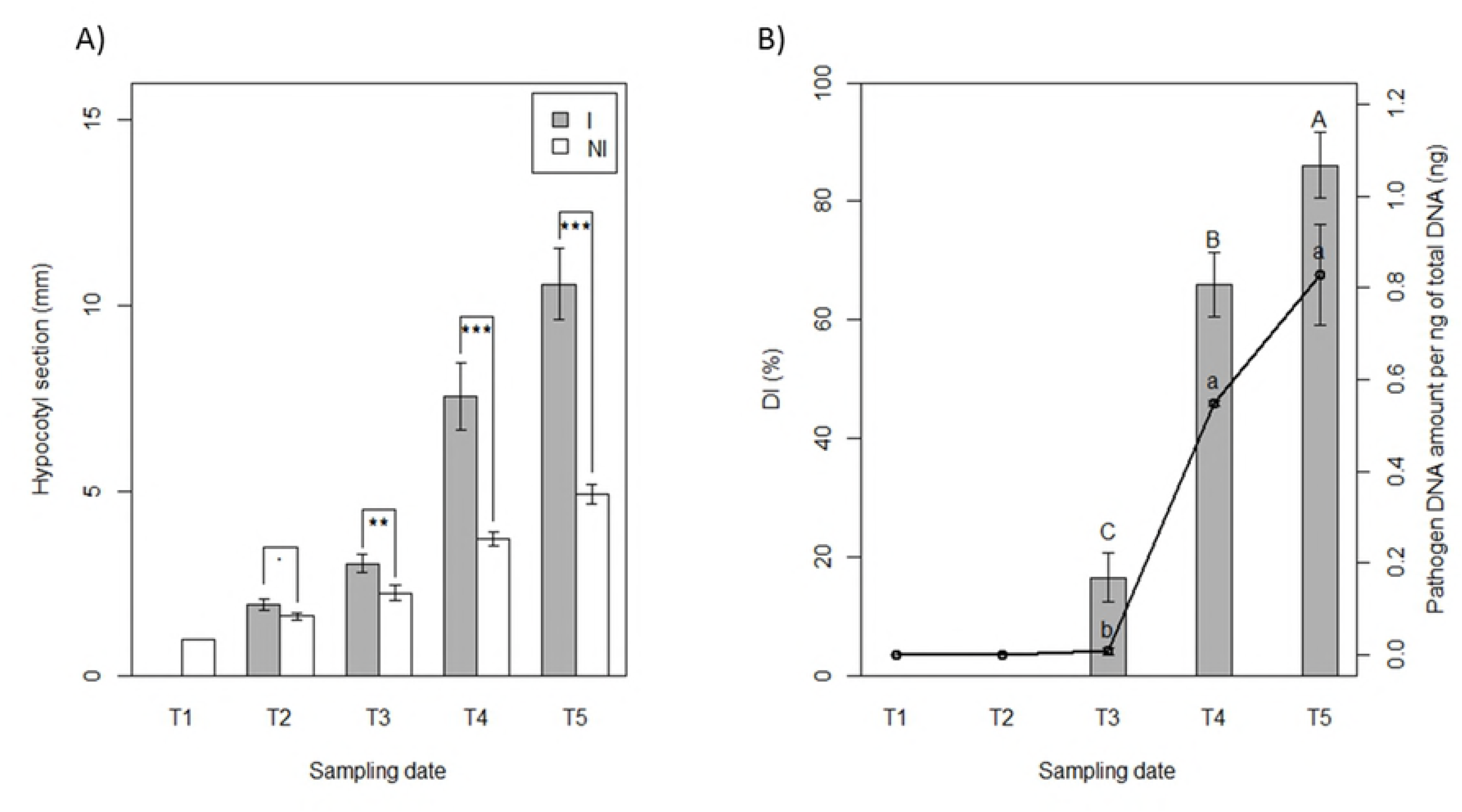
Clubroot disease development. The clubroot development was measured in greenhouse conditions on Chinese cabbage roots over successive sampling at 0 (T1), 7 (T2), 14 (T3), 23 (T4) and 35 (T5) days after inoculation (DAI) by *P. brassicae*. **A)** Taproot width comparison at 1 cm under the soil surface between non-inoculated (NI) and inoculated (I) plants at each date of sampling. Each histogram bar represents the mean of three replicates of four plants. Dot and stars indicate statistically different LSM means (.: P ≤ 0.1, *: P ≤ 0.05, **: P ≤ 0.01, ***: P ≤ 0.001). **B)** Disease index (DI) and pathogen DNA amount (per ng of total DNA) in roots of I plants at each date of sampling. DI indices were represented by the histogram bars and pathogen DNA amount by the black lines. Each dot or histogram bar represents the mean of three replicates of four pooled plants. Capital and lowercase letters indicate statistical differences (p-values ≤ 0.05) between sampling dates for DI and pathogen DNA amounts, respectively.

### Effect of *P. brassicae* on the rhizosphere and root microbiota of Chinese cabbage

#### In the rhizosphere of diseased plants compared to healthy plants

In the rhizosphere compartment, no significant variation of bacterial richness and diversities was measured between healthy and diseased plants whatever the sampling date (T1 to T5) (S5 Fig). The same results were observed in fungal communities except that a reduction of diversity occurred in the rhizosphere of diseased plants at the end of the experiment at T5 (S6 Fig). In bacterial communities, the sampling date explained 21.2% of the overall variance of the data (p = 0.001, 95% CI = 19.5%, 24.3%) and the inoculation condition (I vs NI) 4.4% of this variance. This proportion of the variation, albeit small, was found significant (p = 0.02, 95% CI = 3.3%, 5.3%). The microbial dynamics of healthy and diseased plant communities clearly diverged from T3 to T5 as visualized in PCoA (Fig 3). At T4, there is no variation of bacterial phyla_subphyla relative abundances between healthy and diseased plants (Fig 4), but two OTUs (OTU35 and OTU188) assigned to two genera from the α-Proteobacteria phylum (*Sphingopyxis* and *Rhodobacter*, respectively) and two non-assigned β-Proteobacteria (OTU54 and OTU151) became more abundant in the rhizosphere of diseased than healthy plants (Fig 5). At T5, Proteobacteria (α, β and γ) and Bacteroidetes were consistently more abundant in the rhizosphere of diseased than healthy plants, while both Firmicutes and Acidobacteria were less abundant (Fig 4). At this sampling date, 20 OTUs belonging mainly to the Proteobacteria, Bacteroidetes and Firmicutes phyla were significantly more abundant in inoculated than in non-inoculated plant samples and 8 less abundant (Fig 5). Among these 28 rhizospheric OTUs, the more frequent ones were OTU1 (*Bacillus*) that decreased between T1 and T5 in the rhizosphere of all plants but more drastically in diseased plants especially at T5, and OTU5 (*Flavobacterium*), OTU14 (*Dokdonella*), OTU17 (*Pseudomonas*), OTU35 (*Sphingopyxis*), OTU54 (unknown β-Proteobacteria) which were all significantly more abundant in inoculated than non-inoculated samples at T5 (Fig 6). In fungal communities, the sampling date explained a higher proportion of the variance than in bacterial communities (35%, p = 0.001, 95% CI = 26.1%, 49%), while the inoculation condition (inoculated vs non-inoculated) had no significant effect (3.9%, p = 0.077, 95% CI = 2.8%, 5.6%). Until T4, no variation of fungal phylum frequencies was observed (Fig 4). At T5, while Ascomycota, Basidiomycota and Mucoromycotina were less abundant in the rhizosphere of diseased than healthy plants, no significant variation of Chytridiomycota was observed (Fig 4). However, the major OTU (OTU1) assigned to the Chytridiomycota phylum significantly increased in diseased plant, while four minor OTUs also varied: higher relative abundances for OTU55 and OTU60 but lower for OTU11 and OTU20 in diseased than healthy plant samples at T5 (Fig 7). Higher changes of OTU relative abundances occurred in diseased than healthy plant rhizosphere communities during the time-series experiment (Table 1).

**Fig 3.**
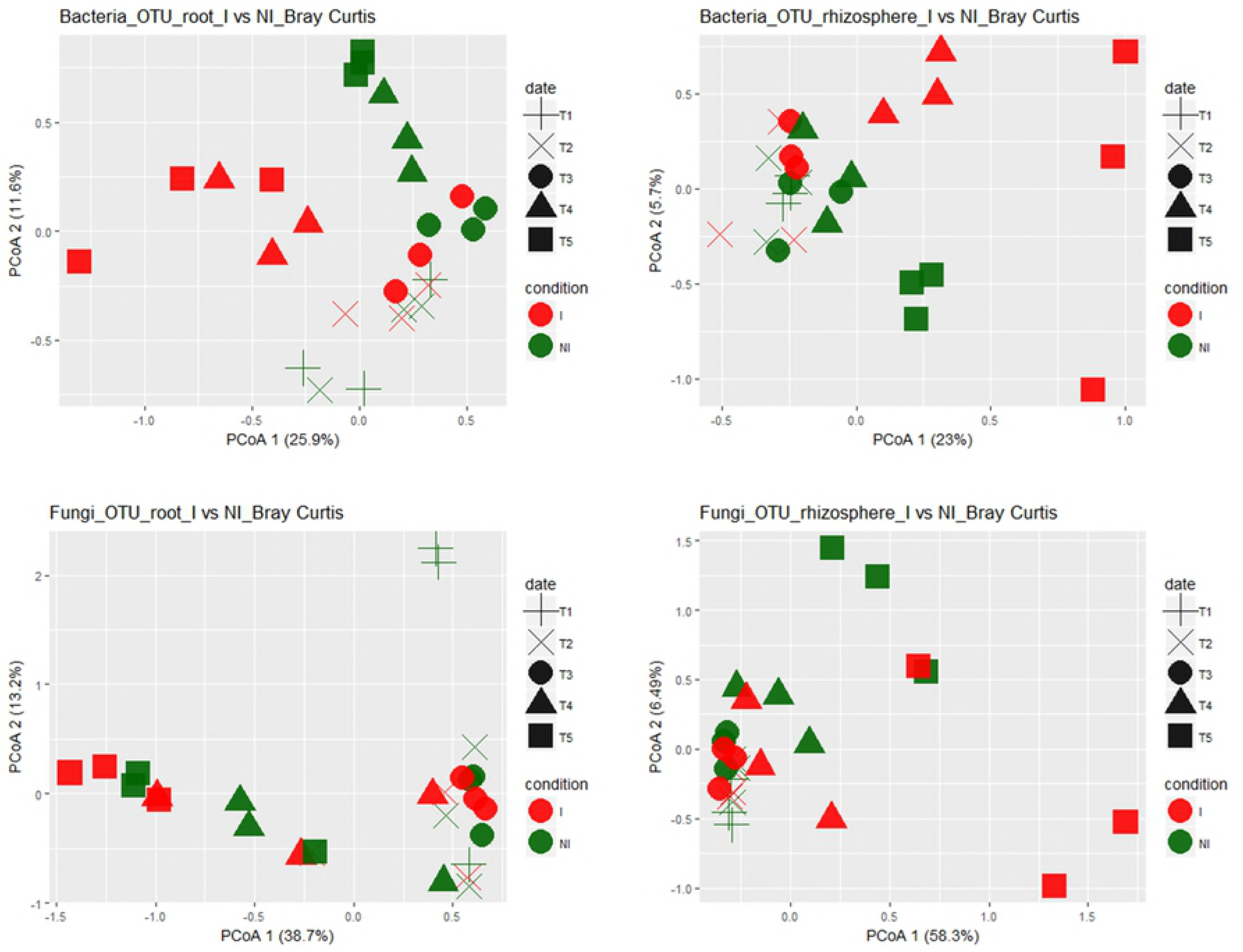
Unconstrained Principal Coordinate Analysis (PcoA) of bacterial and fungal communities from non-inoculated (NI) and inoculated (I) plants. The variances explained by PCoA axes are given in parenthesis. Compartment refers to rhizosphere soil and roots. Sampling date refers to 0 (T1), 7 (T2), 14 (T3), 23 (T4) and 35 (T5) days after inoculation (DAI) with *P. brassicae*, corresponding to plus, crosses, circles, triangles and squares, respectively. Condition refers to non-inoculated (NI) and inoculated (I) plants, represented by green and red colours, respectively.

**Fig 4.**
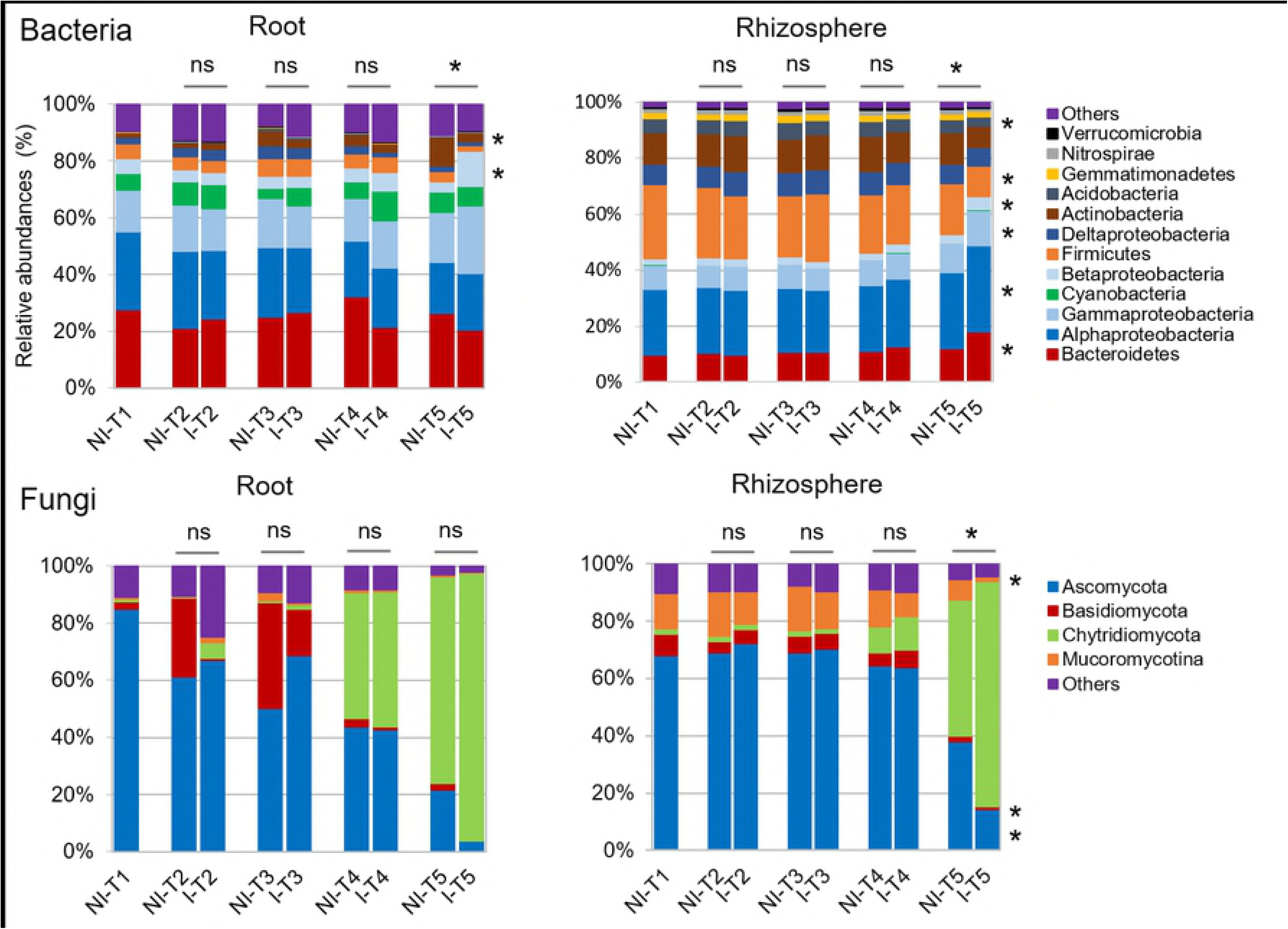
Relative abundances of the most abundant phyla-subphyla in bacterial (A) and fungal (B) communities from root (R) and rhizosphere (RS) compartments. Mean values of abundance (expressed in %) were obtained from three replicates per condition and sampling date. Condition refers to non-inoculated (NI) and inoculated (I) plants. Sampling date refers to 0 (T1), 7 (T2), 14 (T3), 23 (T4) and 35 (T5) days after inoculation (DAI) by *P. brassicae*. Phyla with relative abundances below 1% were grouped as “others”. At each sampling date, significant (p-values ≤ 0.05) and non-significant differences between NI and I plants are indicated by stars and “ns”, respectively. In bacterial communities, the Proteobacteria phylum was divided into four subphyla: α-, β-, γ- and δ-Proteobacteria.

**Fig 5.**
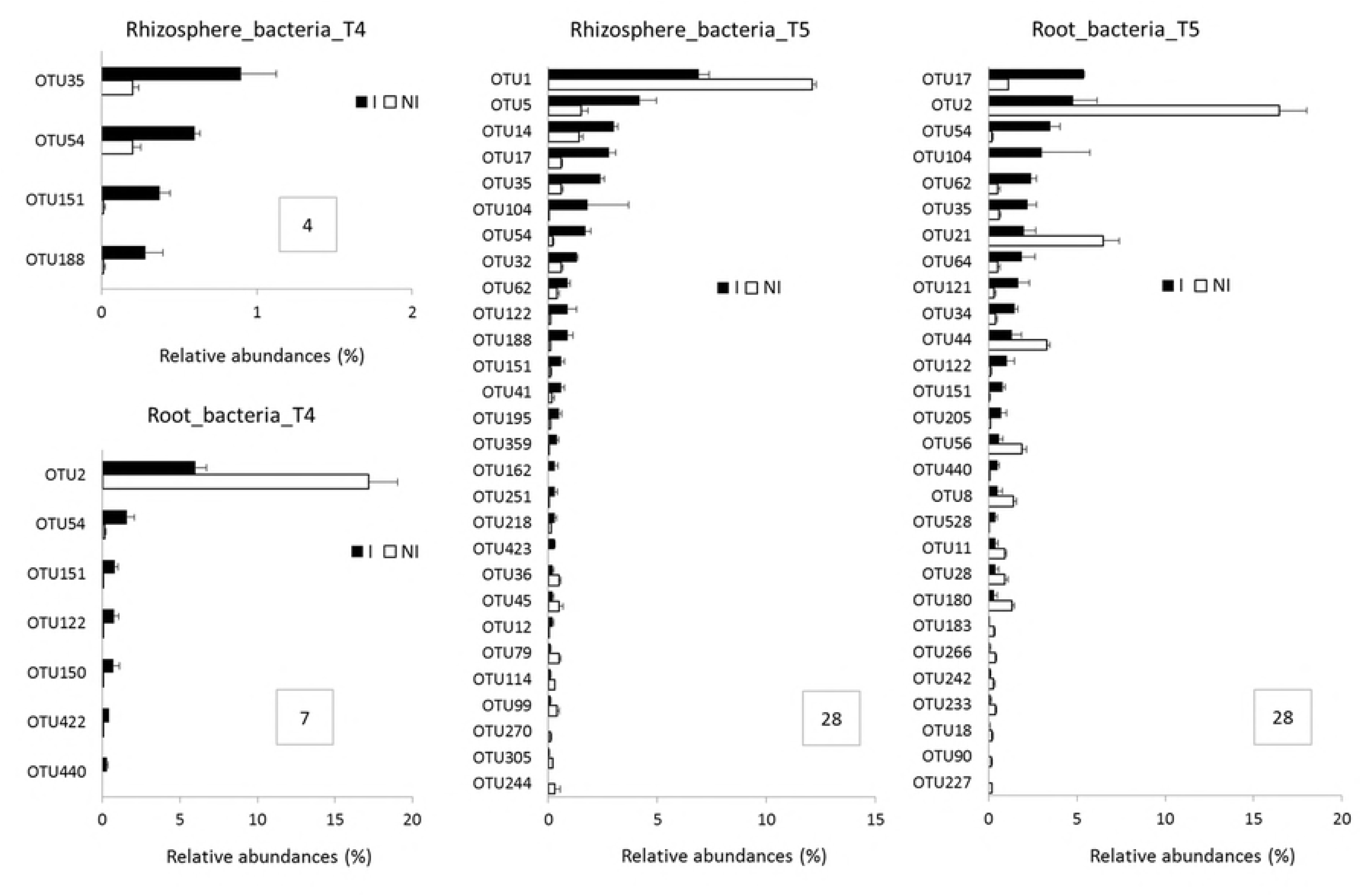
Bacterial OTU relative abundances in the root and rhizosphere microbiota of non-inoculated (NI) and inoculated (I) plants. Bacterial OTUs that significantly differed in their relative abundances (expressed in %) in the root and rhizosphere samples between I and NI plants are represented. Differences were only observed at T4 and T5 in both compartments. Each histogram bar represents the mean RA (± SEM) of three replicates. Only significant differences (p-values ≤ 0.05) between I and NI plants (represented by white and black bars, respectively) are shown, hence at T4 and T5 in both compartments. Framed numbers indicate the number of OTUs with significant different frequencies between I and NI plants.

**Fig 6.**
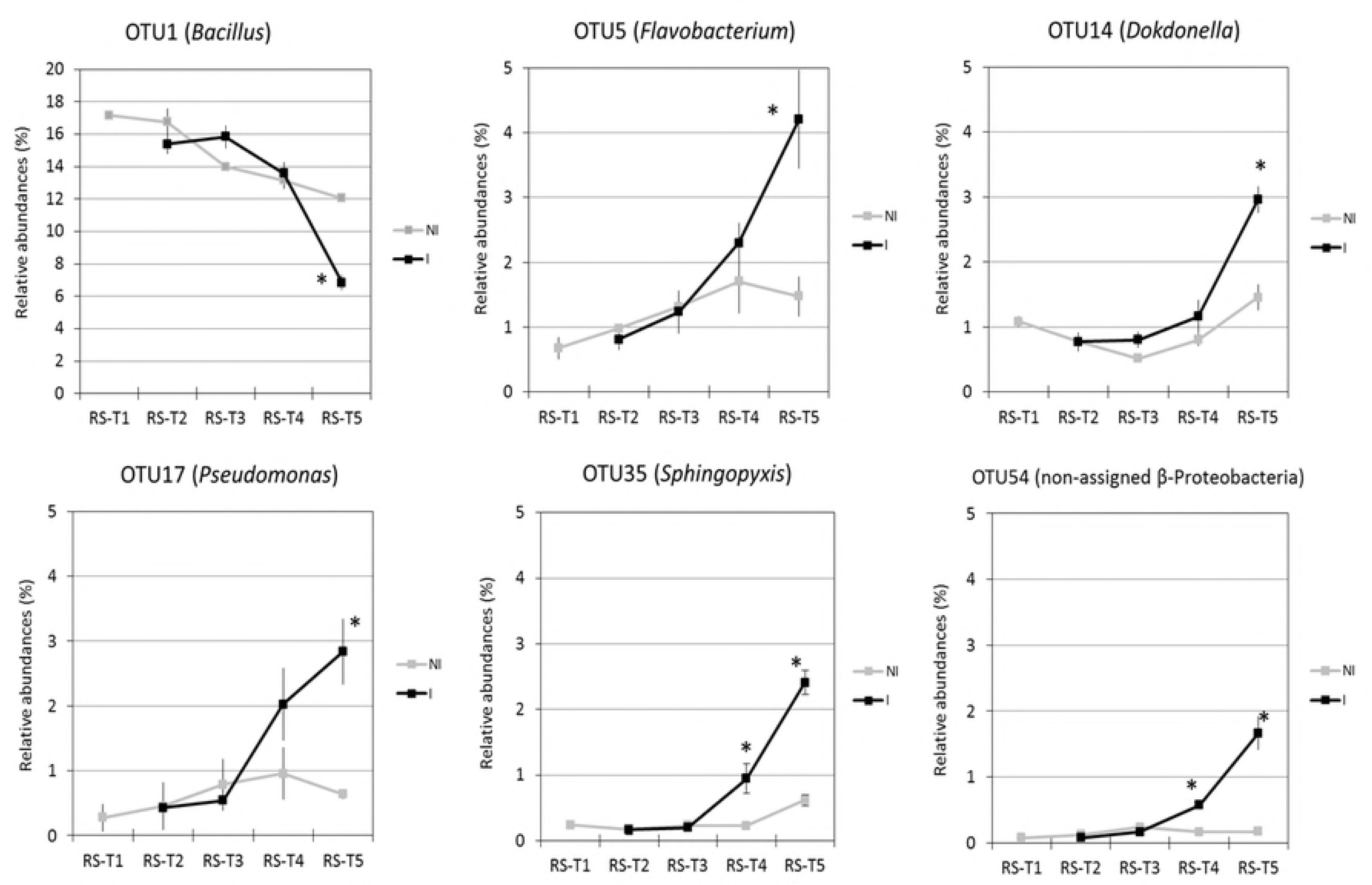
Temporal dynamics of bacterial OTU relative abundances in the bacterial rhizosphere microbiota of non-inoculated (NI) and inoculated (I) plants. Relative abundances (expressed in %) of the most abundant OTUs in rhizosphere (R) at each date of sampling are represented. Sampling date refers to 0 (T1), 7 (T2), 14 (T3), 23 (T4) and 35 (T5) days after inoculation (DAI) with *P. brassicae*. Each dot represents the mean value of relative abundance (± SEM) of three replicates. Stars indicate significant differences (p-values ≤ 0.05) between NI and I plants (represented respectively by black and grey lines) at each sampling date.

**Fig 7.**
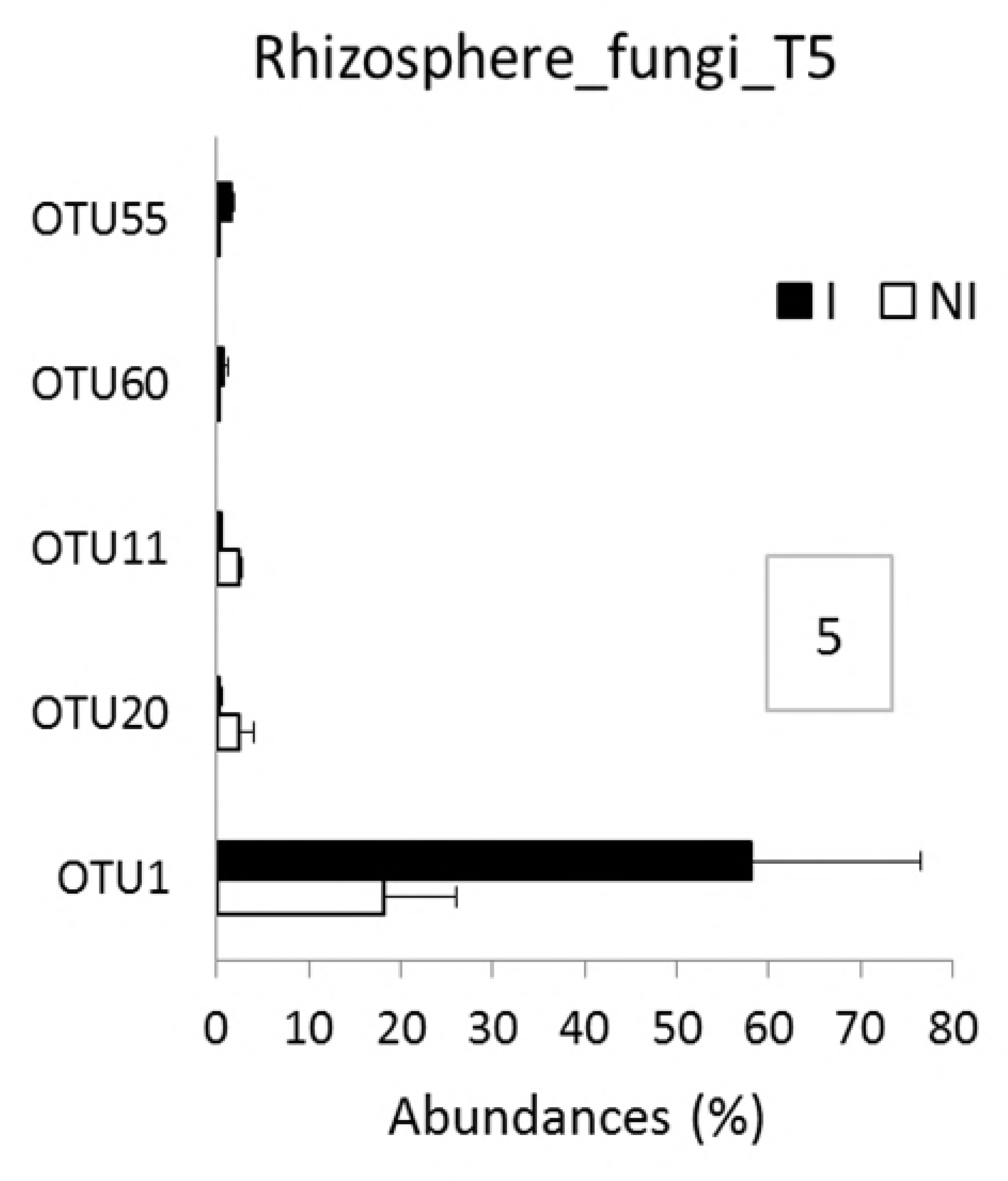
Fungal OTU relative abundances in the rhizosphere microbiota of non-inoculated (NI) and inoculated (I) plants. Fungal OTUs that significantly differ in their relative abundances (expressed in %) in rhizosphere samples (RS) between I and NI plants at T5 are represented. Differences were only observed at T5 in the rhizosphere compartment. Each histogram bar represents the mean relative abundances (± SEM) of three replicates. Only significant differences (p-values ≤ 0.05) between I and NI plants (represented by white and black bars, respectively) are shown, hence at T5 in the rhizosphere compartment. Framed numbers indicate the number of OTUs with significant different frequencies between I and NI plants.

#### Inside the roots of diseased plants compared to healthy plants

In the root compartment, no clear significant differences of bacterial and fungal richness and diversity of communities from healthy and diseased plants were found at each date of sampling (S5 and S6 Figs). In bacterial communities, the sampling date explained 24.4% of the overall data variance (p = 0.001, 95% CI = 20.9%, 28.1%) and the inoculation condition (inoculated vs non-inoculated) 6.2% of this variance (p = 0.002, 95% CI = 4.4%, 8.2%). No significant differences in community composition between inoculated and non-inoculated root samples were observed until T4 when one bacterial OTU (OTU2) assigned to the *Flavisolibacter* genus decreased drastically in relative abundances, while six minor OTUs (OTU54, OTU151, OTU122, OTU150 and OTU422 and OTU440) were slightly more abundant in inoculated than non-inoculated samples (Fig 5). At T5, Actinobacteria were less abundant in the roots of diseased than healthy plants but β-Proteobacteria were more abundant at the phyla-subphyla level (Fig 4). We observed significant differences in relative abundances of 28 OTUs between inoculated and non-inoculated root samples. Among these 28 OTUs, OTU2 (*Flavisolibacter*), OTU21 (*Streptomyces*) and OTU44 (*Pseudomonas*), were the main OTUs which relative abundances had decreased in diseased plants on one hand (Fig 8). On the other hand, the main OTUs which frequencies increased in inoculated vs non-inoculated plants were OTU17 (*Pseudomonas*) but also the two non-assigned β-Proteobacteria OTU54 and OTU62 (Fig 8). Regarding fungal communities, the date of sampling accounted for a higher proportion of the variance than in bacterial communities (36.6%, p = 0.001, 95% CI = 26.2%, 52.9%), while the condition (inoculated vs non-inoculated) had no significant effect (2.7%, p = 0.55, 95% CI = 1.7%, 3.9%) as in the rhizosphere. At each date of sampling, there was no difference in fungal phylum (Fig 4) and OTUs frequencies between diseased and healthy root samples. However, changes of OTU relative abundances occurred in root communities of healthy and diseased plants during the time-series experiment (Table 1).

**Fig 8.**
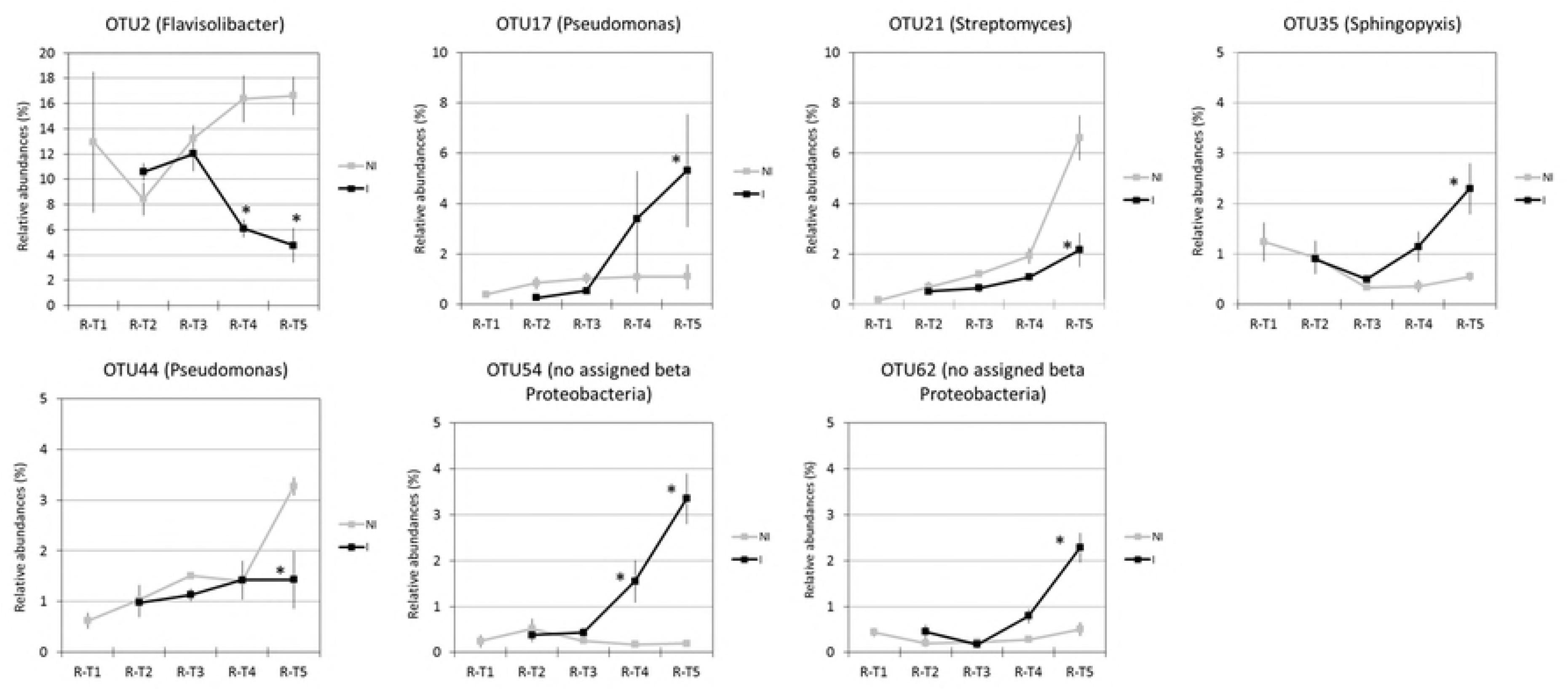
Temporal dynamics of bacterial OTU relative abundances in the root microbiota of non-inoculated (NI) and inoculated (I) plants. Relative abundances (expressed in %) of the most abundant OTUs in root (R) at each date of sampling are represented. Date of sampling refers to 0 (T1), 7 (T2), 14 (T3), 23 (T4) and 35 (T5) days after inoculation (DAI) with *P. brassicae*. Each dot represents the mean value of relative abundance (± SEM) of three replicates. Stars indicate significant differences (p-values ≤ 0.05) between inoculated (I) and non-inoculated (NI) plants (represented by black and grey lines, respectively) at each sampling date.

## Discussion

In our study, the stability of assembled root and rhizosphere communities of Chinese cabbage was investigated by a time-series experiment, during the plant growth and under the effect of the parasitic invasion by *P. brassicae*. During the plant growth, for healthy plants, most of Ascomycota fungi previously recruited by the plant were replaced, mainly in the root compartment, by a Chytridiomycota fungus. The root and rhizosphere-associated community assemblies were also strongly modified by *P. brassicae* infection during the secondary cortical infection stage of clubroot disease.

### A weak but significant rhizosphere effect

Clearly, the communities that assembled in the rhizosphere and bulk soils of healthy plants were very different from the communities found in the roots. These results are consistent with earlier findings on other plant species [2,10,15,59,60]. We found a significant “rhizosphere effect”. Indeed, the alpha diversity analysis showed that the microbiota diversities of the bulk and rhizosphere soils were not distinct from each other. These observations are similar to the findings of several authors in *Arabidopsis thaliana* who reported the resemblance of bacterial communities between rhizosphere and bulk soil samples in multiple soil types [2,10]. At each sampling date, bulk soil and rhizosphere compartments shared a large proportion of OTUs. However, the enrichment of OTUs assigned to the Proteobacteria and Bacteroidetes bacterial phyla, but also to the Chytridiomycota fungal phylum, significantly discriminated rhizosphere from bulk soil samples at the end of the experiment.

### The structure of microbial communities associated with the rhizosphere and roots of healthy plants evolved over time

Roots and rhizosphere of healthy plants were preferentially colonized by Proteobacteria and Bacteroidetes bacterial phyla. Rhizosphere bacterial communities also contained Actinobacteria and Firmicutes but to a higher extent than in the roots. This result was expected because Proteobacteria, Bacteroidetes and Actinobacteria phyla were also highly abundant in the rhizosphere soil of many *Brassicaceae* species like *A. thaliana* [2] and *B. napus* [61-63], with the exception of Bacteroidetes being present at low frequencies in the rhizosphere of *B. napus* cultivated in a Podzol [61] and in a soil collected from an organically managed field [63]. Furthermore, higher frequencies of Firmicutes were observed in rhizosphere communities of Chinese cabbage than in other *Brassicaceae* species. Actinobacteria were detected at lower frequencies in the roots of Chinese cabbage than in the roots of *A. thaliana* [2,10] and *B. napus* [61,62,64]. As in the roots of all Brassicaceae, Cyanobacteria were also abundant in the root of Chinese cabbage.

### A fungus belonging to the Chytridiomycota phylum became dominant in the roots and rhizosphere of non-inoculated plants

The variations of fungal OTU frequencies in the communities of healthy plants were observed mainly at the two last sampling dates. The main changes in fungal communities concerned the relative abundances of an unknown Chytridiomycota which increased drastically at the end of the experiment in the two plant compartments, but especially in roots. This fungus replaced Ascomycota fungi that were previously dominant. In contrast to bacterial 16S sequences, fewer fungal 18S sequences were available to use in taxonomic assignment. However, [65] described the fungal rhizosphere microbiota succession of *B. rapa* plants in compost over three plant generations by sequencing of ITS regions. From the second generation, the Chytridiomycota fungus assigned as *Olpidium brassicae* became dominant in the rhizosphere fungal communities. The organism in our samples could be *O. brassicae* or a close relative, but ultimately this would require confirmation by culturing or more extensive sequencing. This fungus is considered as a soilborne obligate parasite that invades *Brassica* rhizosphere, infects roots and reduces production of pods and seeds [66,67]. Its resting spores can remain dormant in the soil for up to 20 years before infecting roots. However, no symptom was observed on non-inoculated plant roots in our study. For a non-mycorrhized plant, we wondered whether such an endophytic fungus could play a role in plant nutrition or protection against biotic or abiotic stresses as it was observed with *Colletotrichum* spp. [68].

### Clubroot disease altered microbial community structure from the Chinese cabbage roots, then from its rhizosphere

To analyse how the soil borne pathogen affects bacterial and fungal root communities, Chinese cabbage seedlings were inoculated by *P. brassicae* resting spores ten days after sowing. Non-inoculated and inoculated plants were cultivated in controlled conditions for 45 days after sowing. The bacterial and fungal metagenomes from the roots and rhizosphere of heathy and diseased plants were compared at several sampling dates after inoculation. We demonstrated that the invasion by a soilborne parasite changed root and rhizosphere microbial communities already assembled from the soil. Such results about the impact of a soilborne pathogen on the indigenous plant-associated microbiome was also described for *Rhizoctonia solani* on the lettuce microbiome [69] and for *Ralstonia solanacearum* on the tomato rhizosphere microbiota [70].

After inoculation, resting spores of *P. brassicae* released zoospores, which invaded the plant rhizosphere, reached the surface of the root hair and penetrated through the cell wall inside root hairs to form primary plasmodia. After nuclear divisions, the primary plasmodia differentiated into zoosporangia and secondary zoospores were formed in each zoosporangium to be released into the rhizosphere soil [34]. During this primary infection stage, *P. brassicae* was not detected in roots by quantitative PCR indicating that the amount of protist was very low. The interactions between the primary and secondary zoospores and the plant microbiota by direct or indirect mechanisms did not result in detectable changes in bacterial and fungal communities, neither in the roots, nor in the rhizosphere. After being released, the secondary zoospores penetrate the taproot cortical tissues. Inside invaded taproot cells, the pathogen develops into secondary plasmodia which are associated to cellular hypertrophy, followed by gall formation in the tissues [34]. This secondary infection stage was localized inside the roots. During this cortical infection, the amount of *P. brassicae* increased drastically and multiple direct interactions between the protist and the endosphere communities could occur.

We demonstrated that when *P. brassicae* developed inside the roots during its secondary infection stage, it strongly modified the endophytic bacterial communities and lightly the fungal communities. Then, probably as a consequence of the disturbances caused by the interactions between *P. brassicae* and the endophytic communities inside the roots, shifts in rhizosphere communities of diseased plants occurred only at the last date of sampling. Changes in plant microbiota probably occurred by direct microbe-microbe interactions, mainly in the root compartment and then by direct microbial exchanging between the two compartments. However, changes in microbial composition following plant-parasite interactions are often hypothesized to be based on some modifications of the plant chemistry. Salicylic Acid (SA) and Jasmonic Acid (JA) are important hormonal regulators of the plant immune signalling network in which it is commonly accepted that SA is effective against biotrophic and JA against necrotrophic pathogens. However, *P. brassicae* is a biotrophic parasite and both SA and JA signalling pathways could play a role in partial inhibition of clubroot development in compatible interactions between *A. thaliana* and *P. brassicae* [71]. The defence-related phytohormones SA and JA are also known to important modulators of microbiota assembly of *A. thaliana* [72,73]. The accumulation of SA and/or JA or both in plant roots in response to *P. brassicae* infection could lead to modify the composition of plant rhizodeposits and to stimulate specific microbiota in the roots and rhizosphere.

These direct microbe-microbe or indirect microbe-plant interactions could drive the selection of a plant protective microbiome. In few situations, the competitive interaction between soil borne pathogens and root microbiota for available nutrients and microsites could lead to a strong restriction of the pathogen by the activities of specific microorganisms. These situations were already described in suppressive soils for soil borne [74] and foliar parasites [75]. A sequence of events taking place in the rhizosphere of sugar beet seedlings growing in a disease suppressive soil infected by *R. solani* was proposed as a model [74]. The fungus may induce, directly or indirectly via the plant, stress responses in the rhizosphere microbiome by the production of oxalic and phenylacetic acid and lead to shifts in microbiome composition by the activation of Oxalobacteraceae, Burkholderiaceae, Sphingobacteriaceae and Sphingomonadaceae families present in the suppressive rhizosphere microbiome. This stress in turn could trigger a response in these bacterial families, leading to the activation of antagonistic traits that restrict pathogen infection [74].

In light of our results, we also propose a model (Fig 9) in which *P. brassicae* during the first step of its life cycle crosses the plant rhizosphere and infect the root hair without inducing changes in microbiota composition as a consequence of plant metabolism modification. Then, the parasite penetrates inside the roots during the second step of its life cycle and induces due to gall growth strong modifications of the root microbiota. This invasion leads to modification of plant metabolites and root exudates, but also to induction of the plant immune system. As consequences of these trophic and defence modifications, we observed the selection in the root microbiota of specific microorganisms that could (i) use new metabolites, (ii) produce a signal triggering defense responses in plants and (iii) activate, directly or indirectly, other microorganisms in the root and rhizosphere microbiota to control the protist. Future studies will focus on investigating these hypotheses by, among others, selecting other soils and plant genotypes to promote the mechanisms that lead to the restriction of parasitic infection. We also want to develop more functional analysis of the plant-microbiota interactions to identify the underlying mechanisms.

**Fig 9.**
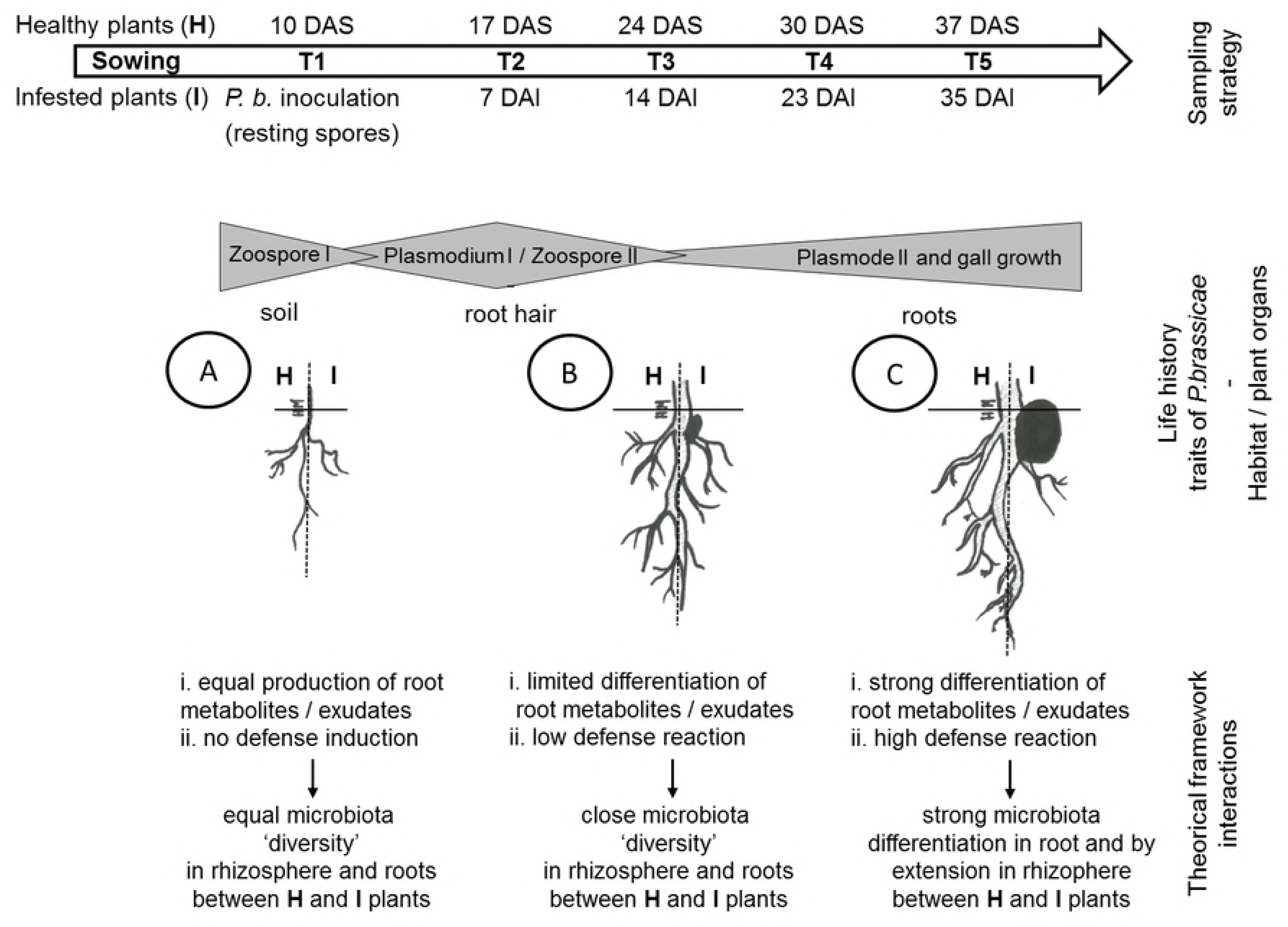
Model illustrating the proposed sequence of events (A to C) taking place in the roots and rhizosphere of Chinese cabbage plant during invasion by *P. brassicae*. Depicted are the changes in microbial community composition in the two compartments as consequences of potential changes in root exudation and plant defense reactions.

The importance of microbiome for the functioning of plant has been widely recognized. Understanding the complex interactions between the pathogen or more generally biotic stress, the plant and its rhizosphere microbiome network are also key elements in shaping a plant-protective microbiome to improve the efficacies of biocontrol agents and partially resistant plants in controlling soil borne plant diseases. By this, plant microbiome is expected to have an important impact in biotechnology and will be a key point for the next Green Revolution as a harbinger to draw a new model for sustainable agriculture.

## Acknowledgment

The authors thank Christine Lariagon and Yannick Lucas for their technical assistance. They also thank Hector Mougel for helping to design to Fig 9 and Maxime Hervé for his advises on statistical analyses. The authors would like to thank the BrACySol biological resource centre (INRA, IGEPP Ploudaniel, France) for providing the seeds used in this study. This work has benefited from the involvement of bioinformatics service of the GenoSol platform from the INRA (French National Institute for Agronomic Research) of Dijon.

## Supporting information

**S1 Fig. Alpha diversity of bacterial communities from non-inoculated (NI) plants.** Richness (i.e. observed OTU) and diversity (i.e. Shannon index) of non-inoculated root (R), rhizosphere (RS) and bulk soil (BS) samples at different sampling dates are represented. Bacterial diversities were estimated with OTUs count data normalized by sample size and rarefied to 1,000 counts. Sampling date refers to 10 (T1), 24 (T3) and 45 (T5) days after sowing (DAS). For each sample, the number of replicates was n = 3. At each sampling date, lowercase letters indicate significant differences (p-values ≤ 0.05) between conditions, which were assessed by ANOVA followed by post hoc Tukey’s HSD test.

**S2 Fig. Alpha diversity of fungal communities from non-inoculated (NI) plants.** Richness (i.e. observed OTU) and diversity (i.e. Shannon index) of non-inoculated root (R), rhizosphere (RS) and bulk soil (BS) samples at different sampling dates are represented. Fungal diversities were estimated with OTUs count data normalized by sample size and rarefied to 1,000 counts. Sampling date refers to 10 (T1), 24 (T3) and 45 (T5) days after sowing (DAS). For each sample, the number of replicates was n = 3. At each sampling date, lowercase letters indicate significant differences (p-values ≤ 0.05) between conditions, which were assessed by ANOVA followed by post hoc Tukey’s HSD test.

**S3 Fig. Unconstrained Principal Coordinate Analysis (PcoA) of the bacterial and fungal communities from non-inoculated Chinese cabbage plants and bulk soil samples**. The variances explained by PCoA axes are given in parenthesis. Compartment refers to bulk soil (BS), rhizosphere soil (RS) and roots (R), represented by orange, brown and green colours, respectively. Sampling date refers to 10 (T1), 24 (T3) and 45 (T5) days after sowing (DAS).

**S4 Fig. Constrained Principal Coordinate Analysis (CPCoA) of the bacterial and fungal communities from non-inoculated Chinese cabbage plants and bulk soil samples**. Compartment refers to bulk soil (BS), rhizosphere soil (RS) and roots (R), represented by orange, brown and green colours, respectively. Sampling date refers to 10 (T1), 24 (T3) and 45 (T5) days after sowing (DAS), represented by crosses, triangles and squares respectively. The variances explained by CPCoA axes are given in parenthesis. For each CPCoA, variations between samples in Bray-Curtis distances were constrained by compartment (in the left column) or sampling date (in the right column) factor. Canonical analysis of principal coordinates (CAP) was performed to quantify the influence of these factors on the β-diversity. The percentage of variation refers to the fraction of the total variance of the data explained by each constrained factor. The p-values indicate if the influence of each of these constrained factors on the β-diversity was significant (p-values ≤ 0.05).

**S5 Fig. Alpha diversity of bacterial communities from inoculated (I) compared to non-inoculated (NI) plants.** Richness (i.e. observed OTU) and diversity (i.e. Shannon index) of root (R) and rhizosphere (RS) samples from NI and I plants were measured at different sampling dates. Bacterial diversity was estimated with OTUs count data normalized by sample size and rarefied to 1,000 counts. Richness and diversity associated NI plants and plants inoculated by *P. brassicae* (I) at each sampling date (T1 to T5) were compared. Sampling date refers to 0 (T1), 7 (T2), 14 (T3), 23 (T4) and 35 (T5) days after inoculation (DAI) with *P. brassicae.* At each sampling date, lowercase letters indicate significant differences (p-values ≤ 0.05) between conditions, which were assessed by ANOVA followed by post hoc Tukey’s HSD test.

**S6 Fig. Alpha diversity of fungal communities from inoculated (I) compared to non-inoculated (NI) plants.** Richness (i.e. observed OTU) and diversity (i.e. Shannon index) of root (R) and rhizosphere (RS) samples from NI and I plants were measured at different sampling dates. Fungal diversity was estimated with OTUs count data normalized by sample size and rarefied to 1,000 counts. Richness and diversity associated to NI and I plants at each sampling date (T1 to T5) were compared. Sampling date refers to 0 (T1), 7 (T2), 14 (T3), 23 (T4) and 35 (T5) days after inoculation (DAI) with *P. brassicae.* At each sampling date, lowercase letters indicate significant differences (p-values ≤ 0.05) between conditions, which were assessed by ANOVA followed by post hoc Tukey’s HSD test.

**S1 Table**. **Comparison of bacterial (B) and fungal (F) OTUs relative abundances in the roots (R) and rhizosphere (RS) of healthy and diseased Chinese cabbage plants**. This table is organized into four tabs corresponding to the description of bacterial OTUs i) from the roots, ii) from the rhizosphere, fungal OTUs iii) from the roots and iv) from the rhizosphere. Mean values of abundance (expressed in %) were obtained from three replicates per condition and sampling date. Condition refers to non-inoculated (NI) and inoculated (I) plants. Sampling date refers to 0 (T1), 7 (T2), 14 (T3), 23 (T4) and 35 (T5) days after inoculation (DAI) by *P. brassicae*. OTUs with relative abundances below 1% were not shown. Significant differences (p-values ≤ 0.05) of OTU frequencies between two samples are indicated by crosses. For example, T2 NI/I refers to the comparison of each OTU frequencies between NI and I plants at T2; NI T1/ I T2 refers to the comparison of each OTU frequencies in communities collected from NI plants at T1 and I plants at T2.

**S2 Table. Quantification of non-inoculated (NI) and inoculated (I) plant traits.** The number of leaves per plant, the shoot and root fresh weight, the plant leaf areas, the plant height and root length were measured during the kinetics of plant growth at 10 (T1), 17 (T2), 24 (T3), 33 (T4) and 45 (T5) days after sowing, corresponding to 0 (T1), 7 (T2), 14 (T3), 23 (T4) and 35 (T5) days after inoculation. At each sampling date, numbers in bold and lowercase letters indicate significant differences (p-values ≤ 0.05) between inoculated (I) and non-inoculated (NI) plants. SEM: standard error of the mean; nd: not determined.

## References

1. Berendsen RL, Pieterse CM, Bakker PA. The rhizosphere microbiome and plant health. Trends Plant Sci. 2012; 17:478–486. doi:10.1016/j.tplants.2012.04.001.

2. Lundberg DS, Lebeis SL, Paredes SH, Yourstone S, Gehring J, Malfatti S, et al. Defining the core *Arabidopsis thaliana* root microbiome. Nature. 2012;488(7409): 86–90. doi:10.1038/nature11237.

3. Vorholt JA. Microbial life in the phyllosphere. Nat Rev Microbiol. 2012;10(12): 828–840. doi:10.1038/nrmicro2910.

4. Vacher C, Hampe A, Porté AJ, Sauer U, Compant S, Morris CE. The phyllosphere: microbial jungle at the plant-climate interface. Annu Rev Ecol Evol Syst. 2016;47: 1–24. doi:10.1146/annurev-ecolsys-121415-032238.

5. Barret M, Briand M, Bonneau S, Préveaux A, Valière S, Bouchez O, et al. Emergence shape the structure of the seed microbiota. Appl Environ Microbiol 2015;81: 1257–66. doi:10.1128/AEM.03722-14.

6. Schiltz S, Gaillard I, Pawlicki-Jullian N, Thiombiano B, Mesnard F, Gontier E. A review: what is the spermosphere and how can it be studied? J Appl Microbiol. 2015;119: 1467–81. doi:10.1111/jam.12946.

7. Berg G, Rybakova D, Grube M, Köberl M. The plant microbiome explored: implications for experimental botany. J Exp Bot. 2016; 67: 995–1002. doi:10.1093/jxb/erv466.

8. Bakker PAHM, Berendsen RL, Doornbos RF, Wintermans PCA, Pieterse CMJ. The rhizosphere revisited: root microbiomics. Front Plant Sci. 2013;4: 165. doi:10.3389/fpls.2013.00165.

9. Mendes R, Garbeva P, Raaijmakers JM. The rhizosphere microbiome: significance of plant beneficial, plant pathogenic, and human pathogenic microorganisms. FEMS Microbiol Rev. 2013;37: 634–663. doi:10.1111/1574-6976.12028.

10. Bulgarelli D, Rott M, Schlaeppi K, Ver Loren van Thermaat E, Ahmadinejad N, et al. Revealing structure and assembly cues for *Arabidopsis* root-inhabiting bacterial microbiota. Nature. 2012;488: 91–95. doi:10.1038/nature11336.

11. Schlaeppi K, Dombrowski N, Garrido-Oter R, Ver Loren van Themaat E, Schulze-Lefert P. Quantitative divergence of the bacterial root microbiota in *Arabidopsis thaliana* relatives. Proc Natl Acad Sci USA. 2014;111: 585–592. doi:10.1073/pnas.1321597111.

12. Lennon JT, Jones SE. Microbial seed banks: the ecological and evolutionary implications of dormancy. Nat Rev Microbiol. 2011;9: 119–130. doi:10.1038/nrmicro2504.

13. Hartmann A, Rothballer M, Schmid M. Lorenz Hiltner, a pioneer in rhizosphere. Microbial ecology and soil bacteriology research. Plant Soil. 2008;312: 7–14. doi:10.1007/s11104-007-9514-z.

14. Hartmann A, Schmid M, van Tuinen D, Berg G. Plant-driven selection of microbes. Plant Soil. 2009;321: 235–257. doi:10.1007/s11104-008-9814-y.

15. Edwards J, Johnson C, Santos-Medellín C, Lurie E, Podishetty NK, Bhatnagar S, et al. Structure, variation, and assembly of the root-associated microbiomes of rice. Proc Natl Acad Sci USA. 2015;112: e911–E920. doi:10.1073/pnas.1414592112.

16. Ranjard L, Dequiedt S, Chemidlin Prévost-Bouré N, Thioulouse J, Saby NP, et al. Turnover of soil bacterial diversity driven by wide-scale environmental heterogeneity. Nature Commun. 2013;4: 1434. doi:10.1038/ncomms2431.

17. Bulgarelli D, Garido-Oter R, Münch PC, Weiman A, Dröge J, Pan Y, et al. Structure and function of the bacterial root microbiota in wild and domesticated barley. Cell Host Microbe. 2015;17: 392–403. doi:10.1016/j.chom.2015.01.011.

18. Mahoney AK, Yin C, Hulbert SH. Community structure, species variation, and potential functions of rhizosphere-associated bacteria of different winter wheat (*Triticum aestivum*) cultivars. Front Plant Sci. 2017;8: 132. doi:10.3389/fpls.2017.00132.

19. Cook RJ, Thomashow LS, Weller DM, Fujimoto D, Mazzola M, Bangera G, et al. Molecular mechanisms of defense by rhizobacteria against root disease. Proc Natl Acad Sci USA. 1995;92: 4197–4201. doi:10.1073/pnas.92.10.4197.

20. Marilley J, Aragno M. Phylogenetic diversity of bacterial communities differing in degree of proximity of Lolium perenne and Trifolium repens roots. Appl Soil Ecol. 1999;13: 127–136. doi:10.1016/S0929-1393(99)00028-1.

21. Bowen GD, Rovira AD. The rhizosphere: the hidden half of the hidden half. In: Waisel Y, Eshel A, Kafkafi U, eds. Plant roots: the hidden half. New York, NY, USA: Marcel Dekker. 1991;641–670.

22. Gransee A, Wittenmayer L. Qualitative and quantitative analysis of water-soluble root exudates in relation to plant species and development. J. Plant Nutr. Soil Sci. 2000;163: 381–385. doi:10.1002/1522-2624(200008)163:4<381::AID-JPLN381>3.0.CO;2-7.

23. Mougel C, Offre P, Ranjard L, Corberand T, Gamalero E, Robin C, Lemanceau P. Dynamic of the genetic structure of bacterial and fungal communities at different development stages of *Medicago truncatula* Gaertn. cv. Jemalong J5. New Phytol. 2006;170: 165–175. doi:10.1111/j.1469-8137.2006.01650.x.

24. Kim B, Song GC, Ryu C-M. Root exudation by aphid leaf infestation recruits root-associated *Paenibacillus* spp. to lead plant insect susceptibility. J. Microbiol. Biotechnol. 2016;26: 549–557. doi:10.4014/jmb.1511.11058.

25. Kong HG, Kim BK, Song GC, Lee S, Ryu C-M. Aboveground whitefly infestation-mediated reshaping of the root microbiota. Front. Microbiol. 2016;7: 1314. doi:10.3389/fmicb.2016.01314.

26. Ourry M, Lebreton L, Chaminade V, Guillerm-Erckelboudt A-Y, Hervé M, Linglin J, et al. Influence of belowground herbivory on the dynamics of root and rhizosphere microbial communities. Front Ecol Evol. 2018;6(91). doi.org/10.3389/fevo.2018.00091.

27. Hardoim PR, van Overbeek LS, Berg G, Pirttilä AM, Compant S, Campisano A, et al. The hidden world within plants: ecological and evolutionary considerations for defining functioning of microbial endophytes. Microbiol Mol Biol Rev. 2015;79: 293–320. doi:10.1128/MMBR.00050-14.

28. Vandenkoornhuyse P, Quaiser A, Duhamel M, Le Van A, Dufresne A. The importance of the microbiome of the plant holobiont. New Phytol. 2015;206: 1196–206. doi:10.1111/nph.13312.

29. Bulgarelli D, Schlaeppi K, Spaepen S, Ver Loren van Thermaat E, Schulze-Lefert P. Structure and functions of the bacterial microbiota of plants. Annu Rev Plant Biol. 2013;64: 807–83. doi:10.1146/annurev-arplant-050312-120106.

30. Weller DM, Raaijmakers JM, Gardener BB, Thomashow LS. Microbial populations responsible for specific soil suppressiveness to plant pathogens. Annu Rev Phytopathol. 2002;40: 309–348. doi:10.1146/annurev.phyto.40.030402.110010.

31. Lugtenberg B, Kamilova F. Plant-growth promoting rhizobacteria. Annu Rev Microbiol. 2009;63: 541–556. doi:10.1146/annurev.micro.62.081307.162918.

32. Raaijmakers JM, Paulitz TC, Steinberg C, Alabouvette C, Moënne-Loccoz Y. The rhizosphere: a playground and battlefield for soilborne pathogens and beneficial microorganisms. Plant Soil. 2009;321: 341–361. doi:10.1007/s11104-008-9568-6.

33. Hwang S-F, Strelkov SE, Feng JIE, Gossen BD, Howard RJ. *Plasmodiophora brassicae*: a review of an emerging pathogen of the Canadian canola (*Brassica napus*) crop. Mol Plant Pathol. 2012;13: 105–113. doi:10.1111/J.1364-3703.2011.00729.x.

34. Kageyama K, Asano T. Life Cycle of *Plasmodiophora brassicae*. J Plant Growth Regul. 2009;28: 203–211. doi:10.1007/s00344-009-9101-z.

35. Fähling M, Graf H, Siemens J. Pathotype separation of *Plasmodiophora brassicae* by the host plant. J Phytopathol. 2003;151: 425–430. doi:10.1046/j.1439-0434.2003.00744.x.

36. Somé A, Manzanares MJ, Laurens F, Baron F, Thomas G, Rouxel F. Variation for virulence on *Brassica napus* L. amongst *Plasmodiophora brassicae* collections from France and derived single-spore isolates. Plant Pathol. 1996;45: 432–439. doi:10.1046/j.1365-3059.1996.d01-155.x.

37. Manzanares-Dauleux MJ, Delourme R, Baron F, Thomas G. Mapping of one major gene and of QTLs involved in resistance to clubroot in *Brassica napus.* Theor Appl Genet. 2000a; 101: 885–891. doi:10.1007/s001220051557.

38. R Core Team. A Language and environment for statistical computing. Available from: https://www.R-project.org/.

39. Bates DM. lme4: Mixed-effects modeling with R. 2010. Available from: http://lme4.r-forge.r-project.org.

40. Lenth RV. Least-squares means: The R package lsmeans. J Stat Softw. 2016;69: 1–33. doi:10.18637/jss.v069.i01.

41. Benjamini Y, Hochberg Y. Controlling the false discovery rate: a practical and powerful approach to multiple testing. J R Stat Soc. 1995;B57: 289–30.

42. Manzanares-Dauleux MJ, Divaret I, Baron F, Thomas G. Evaluation of French *Brassica oleracea* landraces for resistance to *Plasmodiophora brassicae*. Euphytica. 2000b; 113: 211–218. doi:10.1023/A:1003997421340.

43. Hervé M. RVAideMemoire: diverse basic statistical and graphical functions. R package version 0.9-6. 2017. Available from: http://CRAN.R-project.org/package=RVAideMemoire.

44. Plassart P, Terrat S, Thomson B, Griffiths R, Decquiedt S, Lelievre M, et al. Evaluation of the ISO standart 11063 DNA extraction procedure for assessing soil microbial abundance and community structure. PLoS ONE. 2012;7: e44279. doi:10.1371/journal.pone.0044279.

45. Chelius MK, Triplett EW. The diversity of archaea and bacteria in association with the roots of *Zea mays L..* Microb Ecol. 2001;41: 252–263. doi:10.1007/s002480000087.

46. Bodenhausen N, Horton MW, Bergelson J. Bacterial communities associated with the leaves and the roots of *Arabidopsis thaliana*. PLoS One. 2013;8: e56329. doi:10.1371/journal.pone.0056329.

47. Borneman J, Hartin RJ. PCR primers that amplify fungal rRNA genes from environmental samples. Appl Environ Microbiol. 2000;66: 4356–60.

48. LêVan A, Quaiser A, Duhamel M, Michon-Coudouel S, Dufresne A, Vandenkoornhuyse P. Ecophylogeny of the endospheric root fungal microbiome of co-occurring *Agrostis stolonifera.* PeerJ. 2017;5: e3454. doi:10.7717/peerj.3454.

49. Terrat S, Christen R, Dequiedt S, Lelièvre M, Nowak V, Regnier T, et al. Molecular biomass and meta taxogenomic assessment of soil microbial communities as influenced by soil DNA extraction procedure. Microb Biotechnol. 2012;5: 135–141. doi:10.1111/j.1751-7915.2011.00307.x.

50. Terrat S, Dequiedt S, Horrigue W, Lelievre M, Cruaud C, Saby NPA, et al. Improving soil bacterial taxa–area relationships assessment using DNA meta-barcoding. Heredity. 2015;114: 468–475. doi:10.1038/hdy.2014.91.

51. Cole JR, Wang Q, Cardenas E, Fish J, Chai B, Farris RJ, et al. The Ribosomal database project: improved alignments and new tools for rRNA analysis. Nucleic Acids Res. 2009;37: D141–D145. doi:10.1093/nar/gkn879.

52. Quast C, Pruesse E, Yilmaz P, Gerken J, Schweer T, Yarza P, et al. The SILVA ribosomal RNA gene database projects: improved data processing and web-based tools. Nucleic Acids Res. 2013;37: D590–D596. doi:10.1093/nar/gks1219.

53. Oksanen J, Blanchet G, Friendly M, Kindt R, Legendre P, McGlinn D, et al. Vegan: Community Ecology Package. R package version 2.4-3. 2017. Available from: http://CRAN.R-project.org/package=vegan.

54. Anderson MJ, Willis TJ. Canonical analysis of principal coordinates: a useful method of constrained ordination for ecology. Ecology. 2003;84: 511–525. doi:10.1890/0012-9658.

55. Dombrowski N, Schlaeppi K, Agler MT, Hacquard S, Kemen E, Garrido-Oter R, et al. Root microbiota dynamics of perennial *Arabis alpina* are dependent on soil residence time but independent of flowering time. ISME J. 2016;11: 43–55. doi:10.1038/ismej.2016.109.

56. Robinson MD, McCarthy DJ, Smyth GK. edgeR: a Bioconductor package for differential expression analysis of digital gene expression data. Bioinformatics. 2010;26: 139–140. doi:10.1093/bioinformatics/btp616.

57. Anders S, McCarthy DJ, Chen Y, Okoniewski M, Smyth GK, Huber W, et al. Count-based differential expression analysis of RNA sequencing data using R and Bioconductor. Nat Protoc. 2013;8: 1765–1786. doi:10.1038/nprot.2013.099.

58. Robinson MD, Oshlack A. A scaling normalization method for differential expression analysis of RNA-seq data. Genome Biol. 2010;11: R25. doi:10.1186/gb-2010-11-3-r25.

59. de Souza RSC, Okura VK, Armanhi JSL, Jorrín B, Lozano N, da Silva MJ, et al. Unlocking the bacterial and fungal communities assemblages of sugarcane microbiome. Sci Rep. 2016;6: 28774. doi:10.1038/srep28774.

60. Niu B, Paulson JN, Zheng X, Kolter R. Simplified and representative bacterial community of maize roots. Proc Natl Acad Sci USA. 2017;114: E2450–E2459. doi:10.1073/pnas.1616148114.

61. Monreal CM, Zhang J, Koziel S, Vidmar J, Gonzalez M, Matus F, et al. Bacterial community structure associated with the addition of nitrogen and the dynamics of soluble carbon in the rhizosphere of canola (*Brassica napus*) grown in a Podzol. Rhizosphere. 2018;5: 16–25. doi:10.1016/j.rhisph.2017.11.004.

62. Rathore R, Dowling DN, Forristal PD, Spink J, Cotter PD, Bulgarelli D, et al. Crop establishment practices are a driver of the plant microbiota in winter oilseed rape (*Brassica napus*). Front Microbiol. 2017;8: 1489. doi:10.3389/fmicb.2017.01489.

63. Gkarmiri K, Mahmood S, Ekblad A, Alström S, Högberg N, Finlay R. Identifying the active microbiome associated with roots and rhizosphere soil of oilseed rape. Appl Environ Microbiol. 2017;83: e01938–17. doi:10.1128/AEM.01938-17.

64. de Compos SB, Youn J-W, Farina R, Jaenicke S, Jünemann S, Szczepanowski R, et al. Changes in root bacterial communities associated to two different development stages of Canola (*Brassica napus* L. var *oleifera*) evaluated through next-generation sequencing technology. Microb Ecol. 2013;65:593–601. doi:10.1007/s00248-012-0132-9.

65. Tkacz A, Cheema J, Chandra G, Grant A, Poole PS. Stability and succession of the rhizosphere microbiota depends upon plant type and soil composition. ISME J. 2015;9: 2349–2359. doi:10.1038/ismej.2015.41.

66. Hartwright LM, Hunter PJ, Walsh JA. A comparison of *Olpidium* isolates from a range of host plants using internal transcribed spacer sequence analysis and host range studies. Fungal Biol. 2010;114: 26–33. doi:10.1016/j.mycres.2009.09.008.

67. Hilton S, Bennett AJ, Keane G, Bending GD, Chandler D, Stobart R, et al. Impact of shortened crop rotation of oilseed rape on soil and rhizosphere microbial diversity in relation to yield decline. PLoS One. 2013;8: e59859. doi:10.1371/journal.pone.0059859.

68. Hacquard S, Kracher B, Hiruma K, Münch PC, Garrido-Oter R, Thon MR et al. Survival trade-offs in plant roots during colonization by closely related beneficial and pathogenic fungi. Nat Commun. 2016;7: 11362. doi:10.1038/ncomms11362.

69. Erlacher A, Cardinale M, Grosch R, Grube M, Berg G. The impact of the pathogen *Rhizoctonia solani* and its beneficial counterpart *Bacillus amyloliquefaciens* on the indigenous lettuce microbiome. Front Microbiol. 2014;5: 175. doi:10.3389/fmicb.2014.00175.

70. Wei Z, Hu J, Gu Y, Yin S, Xu Y, Jousset A, et al. *Ralstonia solanacearum* pathogen disrupts bacterial rhizosphere microbiome during an invasion. Soil Biol Biochem 2018;118: 8–17. doi:10.1016/j.soilbio.2017.11.012.

71. Lemarié S, Robert-Seilaniantz A, Lariagon C, Lemoine J, Marnet N, Jubault M et al. Both the jasmonic acid and the salicylic acid pathways contribute to resistance to the biotrophic clubroot agent *Plasmodiophora brassicae* in *Arabidopsis*. Plant Cell Physiol. 2015;56: 2158–68. doi:10.1093/pcp/pcv127.

72. Lebeis SL, Paredes SH, Lundberg DS, Breakfield N, Gehring J, McDonald M et al. Salicylic acid modulates colonization of the root microbiome by specific bacterial taxa. Science. 2015;349: 860–4. doi:10.1126/science.aaa8764.

73. Carvalhais LC, Dennis PG, Badri DV, Kidd BN, Vivanco JM, Schenk PM. Linking jasmonic acid signaling, root exudates, and rhizosphere microbiomes. Mol Plant Microbe Interact. 2015; 28: 1049–58. doi:10.1094/MPMI-01-15-0016-R.

74. Chapelle E, Mendes R, Bakker Pahm, Raaijmakers JM. Fungal invasion of the rhizosphere microbiome. ISME J. 2016;10: 265–268. doi:10.1038/ismej.2015.82.

75. Berendsen RL, Vismans G, Yu K, Song Y, de Jonge R, Burgman WP et al. Disease-induced assemblage of a plant-beneficial bacterial consortium. ISME J. 2018;12: 1496–1516. doi:10.1038/s41396-018-0093-1.

